# T4 phage RNA is NAD-capped and alters the NAD-cap epitranscriptome of *Escherichia coli* during infection

**DOI:** 10.1101/2024.04.04.588121

**Authors:** Maik Wolfram-Schauerte, Helene Keuthen, Anastassiya Moskalchuk, Tobias Ueffing, Nadiia Pozhydaieva, Adán Andrés Ramírez Rojas, Daniel Schindler, Stefanie Kaiser, Nicole Pazcia, Katharina Höfer

## Abstract

Nicotinamide adenine dinucleotide (NAD^+^) serves as a cap-like structure on cellular RNAs (NAD-RNAs) across all domains of life, including *Escherichia coli*. Beyond its role in metabolism, NAD^+^ also functions as a regulatory molecule in phage defense mechanisms. However, NAD-RNAs have not yet been identified during bacteriophage infections, and the mechanisms governing their synthesis and degradation in this context remain unknown. To address this gap, we used T4 phage infection of *E. coli* as a defined and well-characterized model system to study NAD-RNAs at the virus-host interface.

Here, we report the first identification and characterization of NAD-RNAs during phage infection. Using time-resolved NAD captureSeq, we identified NAD-capped host and T4 phage transcripts and observed that the set of enriched NAD-capped RNAs varies across different infection phases. Importantly, NAD captureSeq identifies NAD-capped transcripts based on enrichment and does not directly report the fraction of molecules that are NAD-capped. Consequently, temporal changes in enrichment may reflect altered NAD-capping, altered transcript abundance, or a combination of both. We provide evidence that NAD-RNAs are generated by the host RNA polymerase by initiating transcription with NAD^+^ at canonical transcription start sites. By quantifying intracellular NAD^+^ and bulk NAD-capped RNA during infection, we observe parallel decreases in both parameters over the course of infection. Furthermore, we characterize NudE.1, a T4 phage-encoded Nudix hydrolase previously shown to have *in vitro* NAD-RNA decapping activity. Together, our work presents the first time-resolved analysis of an RNA modification in a host–phage system, defining the landscape, dynamics, and turnover of NAD-capped RNAs during infection and providing a framework for future studies addressing their regulatory functions in phage biology.

## Introduction

Bacteriophages are viruses that specifically infect bacteria and represent the most abundant biological entities on earth (Clokie et al. 2011). Due to their presence in virtually all environments inhabited by bacteria, they play a central role in shaping microbial communities and ecosystem dynamics (Bernheim and Sorek 2020). In response to the rapid evolution and high turnover of phage particles, phage–host interactions have become dynamic and multilayered biological systems. This ongoing interplay shapes transcriptional, translational, and metabolic landscapes during infection, reflecting a continuous molecular arms race between phage and host (Ofir and Sorek 2018).

The evolutionary arms race between bacteriophages and their bacterial hosts has led to the emergence of numerous sophisticated bacterial defense systems. These include CRISPR-Cas, abortive infection systems, and mechanisms that interfere with DNA and RNA synthesis (Levy et al. 2015; Bernheim and Sorek 2020; Millman et al. 2020). Phages, in turn, have evolved counter strategies such as anti-CRISPR proteins (Borges et al. 2017). While these defense and counter-defense mechanisms have been studied extensively, much less is known about how phage infection reshapes RNA-centered regulatory layers in both the host and the phage (Pozhydaieva et al. 2024b; Putzeys et al. 2024). In particular, little is known about whether and how RNA modifications contribute to phage–host interactions. Such modifications, collectively referred to as the epitranscriptome, can influence the stability, processing, translation, and molecular interactions of RNA, thereby modulating their function without changing the underlying nucleotide sequence (Schauerte et al. 2021; Wiener and Schwartz 2021; Cappannini et al. 2024). Accordingly, RNA modifications may represent a previously unrecognized regulatory layer in phage–host interactions. Yet, despite the profound transcriptome remodeling that occurs during infection, it is still unknown whether RNA modifications contribute to this process in either bacterial or phage transcripts (Pozhydaieva et al. 2024b).

A well-suited system to address this question is the infection of *Escherichia coli* by bacteriophage T4, one of the best-characterized phage–host model systems, with defined infection stages, extensive molecular tools available and one of the most comprehensively studied bacterial epitranscriptomes (Ouhammouch et al. 1995; Wilkens et al. 1997; Luke et al. 2002; Miller et al. 2003; Nechaev et al. 2004; Geiduschek and Kassavetis 2010; Qi et al. 2015; Guegler and Laub 2021; Wolfram-Schauerte et al. 2022). Phages frequently take over the host’s transcription machinery to ensure efficient expression of their genes, as seen in the recruitment of *E. coli* RNA polymerase (EcRNAP) during T4 phage infection (Koch et al. 1995; Wilkens et al. 1997; Miller et al. 2003). Time-resolved dual transcriptomics has shown that in T4-infected *E. coli*, host transcripts are degraded within minutes of infection, while phage transcripts accumulate in a tightly orchestrated temporal pattern (Luke et al. 2002; Guegler and Laub 2021; Wolfram-Schauerte et al. 2022). These findings indicate that RNA stability is under tight regulation during infection and suggest that mechanisms beyond transcription alone contribute to the dynamic remodeling of host and phage transcriptomes. Thus, in addition to nucleases, RNA secondary structures, and RNA–protein interactions, epitranscriptomic RNA modifications may represent an additional regulatory layer shaping RNA fate during infection. One RNA modification, shown to impair RNA stability in prokaryotes, is the 5′ NAD-cap, in which the ubiquitous redox cofactor nicotinamide adenine dinucleotide (NAD^+^; hereafter referred to as NAD) serves as a non-canonical 5′-terminal cap structure, thereby generating NAD-capped RNAs (Huang 2003; Chen et al. 2009; Cahova et al. 2015; Wang et al. 2019). The 5’-NAD-cap exists in all domains of life, a discovery enabled by the development of specific NAD-RNA sequencing protocols such as NAD captureSeq (Cahova et al. 2015; Winz et al. 2017; Vvedenskaya et al. 2018; Hu et al. 2021; Zhang et al. 2021). In *E. coli*, NAD-capping is mediated by the RNAP, which can use NAD as a non-canonical initiating nucleotide (NCIN) instead of ATP (Bird et al. 2016; Julius and Yuzenkova 2017). This cap protects RNAs from exonucleolytic degradation, whereas decay is initiated by Nudix hydrolases, such as NudC in *E. coli,* which hydrolyze the diphosphate linker in the NAD-cap to produce rapidly degradable 5′-monophosphorylated RNAs (Cahova et al. 2015; Höfer et al. 2016b; Zhang et al. 2016). Together, these findings establish NAD capping and decapping as a mechanistically plausible means to regulate RNA stability.

Because phage infection causes rapid and extensive changes in the abundance and turnover of host and phage transcripts, NAD-(de)capping may represent an important yet unexplored regulatory mechanism in phage–host interactions. However, during phage infections such as T4 phage infection of *E. coli*, the abundance, identity, biosynthesis and turnover of NAD-RNAs remain unknown. It is also unclear whether phage transcripts themselves can be NAD-capped and whether phage-encoded enzymes contribute to NAD-RNA metabolism.

To address this knowledge gap, we performed a time-resolved analysis of the NAD epitranscriptome during T4 infection using an optimized NAD captureSeq workflow combined with long-read sequencing to enable precise 5′-end mapping and full-length transcript identification (Cahova et al. 2015; Winz et al. 2017; Hu et al. 2021; Zhang et al. 2021). This approach allowed us to simultaneously profile NAD-capped host and phage RNAs across the infection cycle. Our time-resolved data reveal infection phase-specific NAD-capping of T4 transcripts alongside a progressive decrease in host NAD-RNA abundance, consistent with the global replacement of host transcripts by phage RNAs during infection. We find that NAD-capped phage RNAs constitute a substantial fraction of phage transcripts and provide evidence that EcRNAP is responsible for NAD capping of both host and phage RNAs across infection stages

Quantifying both NAD and NAD-RNA pool over the time course of infection, we find that NAD-capping and cellular NAD availability may be functionally linked during infection. Moreover, *in vitro*, we show that the T4 Nudix hydrolase NudE.1 hydrolyzes NAD and NAD-RNAs, representing the first phage-encoded decapping enzyme. Its catalytical inactivation delays lysis, underscoring a role of NudE.1 catalytic activity during infection, despite the extensive remodeling of host and phage transcript populations over the course of infection, without detectably altering global NAD or NAD-RNA pools.

Together, this work aims to define the landscape of NAD-capped RNAs during phage infection and to identify enzymatic activities involved in their formation and turnover, thereby providing a foundation for future studies addressing the functional consequences of NAD-capping in phage–host interactions.

## Materials and Methods

### General

All DNA and RNA oligos were purchased from Integrated DNA Technologies. Chemical compounds were obtained from Sigma-Aldrich or Carl Roth GmbH & Co. KG, if not indicated otherwise. RNA was precipitated in the presence of 0.3 M NaOAc pH 5.5 and 1 volume isopropanol by centrifugation at 21,000 x *g* or 18,500 x *g*, 4 °C for 90 min after incubation at −20 °C overnight, if not indicated differently.

### Strains, media and cultivation

Cultivation of *E. coli* strains (Supplementary Table S1) was conducted in lysogeny broth (LB) medium supplemented with the 30 µg/µL kanamycin (for pET28a-based expression strains) or without antibiotic (*E. coli* B strain) at 37 °C, 180 rpm.

### Determination of *E. coli* cell concentration

*E. coli* cell concentration was determined by diluting an *E. coli* culture at an optical density at 600 nm (OD_600_) = 0.84 1:2,000 in isotone (Sigma-Aldrich) and measuring 75 µL thereof at a Coulter counter (Multisizer 4e (Beckman Coulter)). Cells counts in the range of 1 – 2 µm particle size were summed and blank background was subtracted to obtain the measure of 3 * 10^8^ *E. coli* cells per mL at OD_600_ = 0.8.

### T4 phage infection

Bacteriophage T4 infection of *E. coli* strain B was performed in LB medium, supplemented with 1 mM CaCl_2_ and 1 mM MgCl_2_, either at room temperature (rt) shaking at 120 rpm or at 37 °C shaking at 180 rpm. OD_600_ of *E. coli* culture used for infection varied between 0.2 – 0.8 and phages were added to a final multiplicity of infection (MOI) ranging from 0.2 – 5.0.

### T4 phage production

*E. coli* BL21 DE3 pET28a was grown to OD_600_ = 0.6 in LB medium with kanamycin at 37 °C, 180 rpm. T4 phage solution was added to a multiplicity of infection (MOI) of 0.2 and the culture incubated at room temperature, 120 rpm overnight. Unlysed cells were pelleted by centrifugation at 5,000 x *g*, 4 °C for 25 min and supernatant was filtered through 0.22 µm filters. The resulting T4 phage solution was stored in glass bottles at 4 °C.

### Determination of T4 phage titer via plaque assay

T4 phage solution was diluted 10^-1^ to 10^-11^ in LB medium. 300 µL of *E. coli* B strain bacterial culture (OD_600_ = 1.0) and 100 μL of respective diluted phage solution were mixed each and incubated for 2-3 min at rt. Then, samples were mixed each with 4 mL of soft agar (2 % w/v agar, 1.5 % w/v pepton, 0.75 % w/v yeast extract, 1.5 % w/v NaCl) preheated to 55 °C. Whole mixtures were poured onto LB agar plates and incubated at 37 °C overnight. The titer was determined using equation (1):

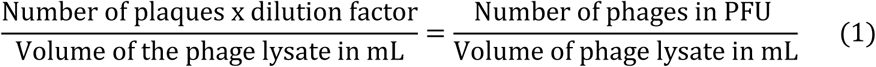

*E. coli* B strain was grown at 37 °C, 180 rpm in LB medium supplemented with 1 mM CaCl_2_ and 1 mM MgCl_2_ to OD_600_ = 0.8. Then, cultures were transferred to rt shaking at 120 rpm and phage solution was added to an MOI of 1.5 – 5.0. OD_600_ was regularly monitored to determine cell lysis. Alternatively, the same assay was performed in a BioTek plate reader (Agilent Technologies) measuring cell density as OD_600_ in a 48-well format at rt shaking at 300 rpm every 3 min. For the plate reader measurement of OD_600_, OD values were normalized to 1 at time point 0 before plotting.

### Burst size assay

50 mL of *E. coli* B strain culture in LB medium supplemented with 1 mM CaCl_2_ and 1 mM MgCl_2_ that was grown to OD_600_ 0.8 was infected with T4 phage (WT or NudE.1 E64,65Q) at MOI 0.01 and incubated in a water bath at 37 °C. To determine the amount of all phages added, 50 µL of appropriately diluted culture at 6 minutes post infection was subjected to plaque assay as described above. At the same time point and all other subsequent time points, the culture was diluted according to the expected number of phages (that can be assessed by plaque assay) and 1 mL free-floating phages were specifically isolated with 50 µL chloroform. 50 µL thereof were used for plaque assay as described above. The amount of initially infecting phages was calculated as the difference between all phages and free-floating phages at 6 min post infection. This was used to calculate the burst size during the first 60 min of infection.

### T4 phage mutant generation and screening

*E. coli* BL21 DE3 containing plasmids pET28a_NgTET_NudE.1-E64,65Q and pCpf1_NudE.1 was used for mutagenesis. The cells were grown to OD_600_ = 0.4 at 37 °C, 160 rpm, when NgTET expression was induced by addition of 50 µM IPTG and cells were grown further to OD_600_ = 0.8. The cultivation was proceeded until OD_600_ of 0.8. At this point, the cultures were adjusted to room temperature shaking at 120 rpm and supplemented with 1 mM MgCl_2_ and 1 mM CaCl_2_. Infection was carried out with NgTET-treated T4 phage WT set to a multiplicity of infection (MOI) of 0.5 as described previously (Pozhydaieva et al. 2024a). The infection was performed for 3 h, at 130 rpm and 23 °C after which the cells were pelleted and supernatant filtered through 0.45 µm filters. The titer of the newly generated phages was determined via plaque assay and the phages were used for the counter-selection via infection of *E. coli* Cas13a_NudE.1 (plasmid pBA560-Cas13a-NudE.1). The counter-selection was performed under the same conditions as the mutagenesis. The counter-selected phages were filtered through 0.45 µm filter and used for a plaque assay with *E. coli* B strain. 144 individual plaques were picked and resuspended in 200 µL Mg-Pi-buffer (26 mM Na_2_PHO_4_, 68 mM NaCl, 22 mM KH_2_PO_4_, 1 mM MgSO_4_, pH 7.5) supplemented with 2 µL Chloroform and incubated at room temperature shaking at 500 rpm for 1 h. Plaques in solution were stored at 4 °C. The genotype of individual phage isolates was subsequently verified by long-read amplicon sequencing. Phages were screened for mutations by Nanopore sequencing as described previously (Pozhydaieva et al. 2024a; Ramirez Rojas et al. 2024). 1 µL resuspended plaque were used for each PCR in 10 µL scale in the presence of 1 x High Fidelity PCR Master Mix (NEB) and 500 nM NudE.1 fwd and rev screening primer (Supplementary Table 2) using 98 °C for 3 min followed by 30 cycles of 98 °C for 10 s, 60 °C for 10 s and 72 °C for 1 min with a final hold at 72 °C 10 min. A second dual barcoding PCR was performed using KAPA HiFi HotStart ReadyMix (Roche) in the presence of 1 µL of a 1:10 dilution of the initial PCR as template and 0.3 µM of the barcoding primers in 7 µL reactions. PCRs were conducted at 95 °C for 3 min followed by 20 cycles of 98 °C for 20 s, 66 °C for 15 s and 72 °C for 60 s with a final extension of 72 °C for 5 min. All the barcoded PCR reactions were pooled and purified using an open-source purification procedure using the NucleoMag kit for NGS library preps (Macherey Nagel) (Oberacker et al. 2019). Briefly, DNA was bound to magnetic beads, washed twice with 80 % ethanol and eluted in 100 µL elution buffer (5 mM Tris-HCl pH 8.5). Concentration was determined with NanoDrop (Thermo Fisher Scientific) and Qubit (Invitrogen) using the broad range and/or high sensitivity assay. Sequencing libraries were generated with the SQK-LSK109 Ligation Sequencing kit (Oxford Nanopore Technologies) according to the manufacturer’s guidelines starting with 1 µg of input DNA. Sequencing was performed on Flongle flow cells (R9.4.1 chemistry) using a MinION device. The resulting Nanopore sequencing data were subsequently processed and analyzed as follows. Raw reads were basecalled using Guppy (v6.4.2). Reads were analysed using the CRISPRT4 pipeline (Pozhydaieva et al. 2024a) (https://github.com/MaikTungsten/CRISPRT4). In a miniconda environment, reads were demultiplexed using minibar (Krehenwinkel et al. 2019) and mapped to the T4 phage reference genome (NCBI: NC_000866.4, accessed 15^th^ September 2023) using minimap2 (version 2.24) (Li 2018). The resulting SAM files were converted to BAM files, sorted, and indexed with samtools (version 1.4.1) (Danecek et al. 2021). Variant calling in the target region of mutagenesis was subsequently performed using longshot (version 0.4.1) (Edge and Bansal 2019) and resulting VCF files were inspected for desired point mutants with Integrative Genomics Viewer (IGV 2.16.0) (Thorvaldsdottir et al. 2013).

Candidate mutant phages carrying the desired mutations were subsequently propagated by adding 10 µL of resuspended plaque material to 10 mL of *E. coli* B culture in LB medium supplemented with 1 mM MgCl₂ and 1 mM CaCl₂, followed by overnight incubation at room temperature and 120 rpm. Sanger sequencing was performed to validate the mutation. As a small fraction of the phage population contained the WT locus of NudE.1, the phage mutant candidate was counter-selected against WT. Therefore, 10 mL *E. coli* BL21 DE3 with plasmid pBA560-Cas13a-NudE.1 was grown in LB medium supplemented with 30 µg/mL chloramphenicol to OD_600_ = 0.4, when the expression of the Cas13a system was induced by addition of tetracycline to final concentration 1 nM. After 10 min incubation, 10 µL phage was added and incubated at 37 °C for 1 h. Thereafter, phage was purified by filtration through 0.45 µm filters, diluted in LB medium and used for plaque assay with *E. coli* BL21 DE3 with plasmid pET28-NudE.1 induced with 1 mM IPTG. Single plaques were picked again, screened with Sanger sequencing and mutant phage amplified as described above.

### Total RNA isolation

100 mL LB medium with 1 mM MgCl_2_ and 1 mM CaCl_2_ were inoculated with overnight culture of *E. coli* strain B, *E. coli* JM109 + pUC19 (with ampicillin as an antibiotic) OD_600_ = 0.1. The culture was incubated at 37 °C, 160 rpm to OD_600_= 0.8. Then, the culture was transferred to rt and the first 5 mL sample was taken (0 min). Subsequently, the bacterial culture was infected with T4 phage solution to a multiplicity of infection of 1.5 or 5.0, respectively. The T4 phage-infected culture was incubated at rt, 120 rpm and 5 mL samples were taken at 1, 4, 7, 10 and 20 min of post-infection or as indicated. After sampling, cells were lysed immediately using one volume (5 mL) of the 90 °C lysis solution (2 % SDS, 4 mM EDTA) and incubated at 90 °C for 2 minutes. RNA was extracted in one volume (10 mL) of Roti-Aqua-Phenol (Carl Roth) by incubating at 67 °C for 10 min and centrifuging for 10 min and at 18,500 x *g*, rt. RNA was extracted again from the upper phase using one volume (10 mL) of Roti-Aqua-P/C/I (Carl Roth) and samples were centrifuged at 18,500 x *g*, 4 °C for 10 min. Upper phases were mixed with one volume of ice-cold isopropanol and 0.1 x volume of 3 M NaOAc pH 5.5 and incubated at −20 °C overnight. RNAs were precipitated by centrifugation at 18,500 x *g* at 4 °C for 2 h. RNA-pellets were resuspended in 450 μL RNase-free water. RNA was digested in 1 x DNaseI buffer with 0.8 U/μL of DNaseI (Roche) to remove residual DNA. Reactions were incubated at 37 °C for 40 min. DNaseI digestion was stopped by addition of one volume of Roti-Aqua-P/C/I. Samples were centrifuged at 17,000 x *g*, 1 min, rt. P/C/I-extraction was performed twice. Residual phenol was removed by diethylether extraction using one volume diethylether and phase separation by short centrifugation. The extraction was performed three times. The residual diethylether was evaporated using the SpeedVac (Thermo Fisher Scientific). RNAs were ispropanol precipitated and RNA pellets were resuspended in 20 – 100 µL RNase-free water. Final RNA concentrations were determined using the NanoDrop (Thermo Fisher Scientific). RNAs were analyzed by 10 % PAGE as well as 1 % agarose gel electrophoresis. Gels were imaged at ChemiDoc (Bio-Rad) using either SYBR Gold (Invitrogen) or PeqGreen (PeqLab) staining. Further, RNAs were analyzed using RNA 6000 Nano Kit and the Bioanalyzer system (Agilent Technologies). Thereby, RNA integrity was determined and only RNAs with RNA integrity numbers (RINs) greater than 9.0 were used for further analysis.

### Preparation and characterization of NAD-RNAs by *in vitro* transcription

For the preparation of Qβ and RNAI transcripts by *in vitro* transcription, the corresponding dsDNA templates were first amplified by PCR. Amplification was performed using 50 nM forward and reverse ultramers (Supplementary Table S2) in the presence of 1 × GC buffer, 5% DMSO, 0.1 mM dNTP mix, 500 nM forward primer, 500 nM reverse primer, and 0.01 U/µL Phusion polymerase (Thermo Fisher Scientific). DNA concentrations were measured using a NanoDrop spectrophotometer, and the amplified templates were subsequently used for *in vitro* transcription. *In vitro* transcription (IVT) of 5′-triphosphate (PPP) Qβ-RNA and RNAI was performed in a 20 µL reaction using the HighYield T7 RNA Synthesis Kit (Jena Bioscience) according to the manufacturer’s instructions in the presence of 1 µg dsDNA template. NAD-capped RNA was generated under the same conditions except for the addition of 3.75 mM ATP and 7.5 mM NAD. The IVT reactions were incubated at 37 °C for 3 h. Residual DNA was removed by DNase I digestion using 20 U DNase I and 1 x DNase I incubation buffer and incubating the mixture at 37 °C for 30 min. *In vitro* transcribed 5’-PPP-RNAs and 5’-NAD-RNAs were purified by 10 % preparative PAGE. RNA bands were visualized by UV shadowing. The bands were excised, and RNA was eluted from the gel material by incubation in 4 mL 0.3 M NaOAc pH 5.5 by shaking at 600 rpm, 14 °C overnight. An additional elution step was performed for 3 h at the same settings. After removal of gel pieces, RNA was isopropanol precipitated, RNA pellets were air-dried and resuspended in 200 μL RNase-free water, isopropanol-precipitated again and resuspended in a final volume of 50 µL RNase-free water. RNA concentrations were measured on the NanoDrop. NAD-RNAs were analyzed using 10 % PAGE. The presence of the NAD-cap was confirmed by 6 % APB-PAGE and NudC (self-purified) digest as described before (Nübel et al. 2017) except for the usage of a HEPES buffered system. Reactions were prepared for analysis in a 10 μL scale by incubating 100 ng of each RNA, 1 x degradation buffer (12.5 mM Tris-HCl pH 7.5, 25 mM NaCl, 25 mM KCl, 5 mM MgCl_2_) and 4.2 μM NudC (1 µL) or 1 μL RNase-free water, respectively. The reactions were incubated at 37 °C, 30 min and applied to 6 % (w.r.t. final acrylamide concentration) APB-PAGE. Gels were imaged at ChemiDoc after staining in SYBR Gold (Invitrogen).

### NAD captureSeq library prep

NAD captureSeq was performed in a similar manner as described in (Winz et al. 2017). To selectively label NAD-capped RNAs, total RNA was subjected to a two-step chemo-enzymatic labeling procedure consisting of an ADPRC reaction followed by a SPAAC reaction. ADPRC reactions were performed in a 100 μL volume containing 1 × ADPRC buffer, 10% 3-azido-1-propanol, 70 μg total RNA from *E. coli* (before and after 1, 4, 7, 10, and 20 min of infection), 10 ng of a 100 nt NAD-RNA spike-in control (Supplementary Table S3), and either the presence or absence of 0.85 μM ADPRC. The mixtures were incubated at 37 °C for 45 min. Reactions were stopped by Roti-Aqua-PCI extraction, followed by diethylether extraction. Then, reactions were isopropanol-precipitated in the presence of 40 µg RNA-grade glycogen. Pellets were resuspended in 20.3 μL RNase-free water. SPAAC reactions were performed in a 40 μL scale in the presence of 20 μL RNA (∼ 70 μg), 1 x PBS and 0.25 mM biotin-PEG4-DBCO (in DMSO). Reactions were incubated at 37 °C, 2 h. Then, 160 μL RNase-free water were added. Reactions were stopped by Roti-Aqua-PCI extraction, followed by diethylether extraction Samples were isopropanol-precipitated in the presence of 40 µg RNA-grade glycogen. RNA pellets were resuspended in 40.3 μL Immobilization buffer (10 mM Na-HEPES pH 7.2, 1 M NaCl, 5 mM EDTA).

The integrity of RNAs after both ADPRC and SPAAC reactions was assessed by agarose gel electrophoresis using 0.3 μL (∼1 μg) of each sample. The labeled RNAs were subsequently subjected to streptavidin-based enrichment. Mobicol classic columns (MoBiTec) were assembled with small filters and placed into 2 mL reaction tubes, and 50 μL streptavidin Sepharose was added to each column. The streptavidin beads were equilibrated by washing three times with 200 μL immobilization buffer. The columns were then centrifuged at 17,000 x *g*, 1 min, rt, and the supernatants were discarded. All further wash steps mentioned were performed three times with the same volume of respective buffer with an exception for washing with streptavidin wash buffer (8 M urea, 50 mM Tris-HCl pH 7.4) and 0.25 x streptavidin wash buffer (2 M urea, 50 mM Tris-HCl pH 7.4). In these cases, the beads were washed five times. After washing, the beads were blocked by addition of 100 μL immobilization buffer, supplemented with 100 μg/mL acetylated BSA and by incubation at 20 °C, 1,000 rpm for 20 min. Then, blocking reactions were centrifuged at 17,000 x *g*, rt for 1 min and flow-through was discarded. Blocking steps were always performed in the same manner with 100 μL of respective buffer and 100 μg/mL acetylated BSA. After blocking, the streptavidin beads were washed with immobilization buffer as previously described. The biotinylated RNAs were captured by streptavidin beads by addition of their complete volume of 40 μL (∼ 70 μg) to each column and incubation at 20 °C, 1,000 rpm for 1 h. The mixtures were centrifuged at 17,000 x *g*, rt for 1 min and the supernatants were discarded. The beads were subsequently washed with streptavidin wash buffer.

Captured RNAs were then subjected to 3′ adaptor ligation to facilitate downstream reverse transcription and sequencing library preparation. The streptavidin beads were equilibrated, blocked, and washed as described above using 1 × standard ligation buffer (500 mM Tris-HCl pH 7.4, 100 mM MgCl₂). The adaptor ligation mixture consisted of 5 µM adenylated 3′ adaptor, 1 × standard ligation buffer, 15% DMSO, 50 μg/mL adenylated BSA, 50 mM 2-mercaptoethanol, 0.5 U/µL T4 RNA ligase 1 (NEB), and 10 U/µL T4 RNA ligase 2 truncated K227Q (NEB). 30 µL of this mixture were added to the beads and the columns were incubated at 4 °C overnight. Then, NaCl was added to the beads to a final concentration of 1.5 M and the columns were incubated at 20 °C, 1,000 rpm for 1 h.

After incubation, reactions were centrifuged at 17,000 × *g* for 1 min at room temperature. The supernatants were discarded, and the beads were washed with streptavidin wash buffer. To convert the captured and adaptor-ligated RNAs into cDNA for subsequent amplification and sequencing, reverse transcription was performed directly on the streptavidin-bound RNA. For this purpose, the beads were equilibrated, blocked, and washed as described above using 1 × first-strand buffer (250 mM Tris-HCl pH 8.3, 375 mM KCl, 15 mM MgCl₂).

For the reverse transcription, the reaction mixture containing 5 μM RT primer, 0.5 mM dNTP mix, 50 μg/mL adenylated BSA, 5 mM DTT, 1 x first-strand buffer and 10 U/µL Superscript IV reverse transcriptase (Thermo Fisher Scientific) was prepared on ice. 30 μL of this mixture were added to the beads and the reactions were incubated at 40 °C for 1 h. Then, dissociated hybrids of biotin–RNA and cDNA were rebound by addition of NaCl to a final concentration of 1.5 M and incubation at 20 °C, 1,000 rpm for 1 h. Thereafter, reactions were centrifuged at 17,000 x *g*, rt for 1 min.

The supernatants were discarded, and the beads were washed with 0.25 × streptavidin wash buffer. Residual free primers were subsequently removed by Exonuclease I digestion. To this end, the streptavidin beads were equilibrated, blocked, and washed as described above using 1 × Exonuclease I buffer (670 mM glycine-KOH pH 9.5, 67 mM MgCl₂, 100 mM 2-mercaptoethanol). For the ExoI digest, the ExoI reaction mixture containing 1 x ExoI buffer and 1 U/μL ExoI (NEB) enzyme was prepared. 30 μL of this mixture were added to the beads and the reactions were incubated at 37 °C for 30 min. After 30 min of incubation, 1.5 μL of ExoI enzyme were added and reactions were incubated for another 20 min. Then, reactions were centrifuged at 17,000 x *g*, rt for 1 min.

The supernatants were discarded, and the beads were washed with 0.25 × streptavidin wash buffer. To enable downstream PCR amplification, cDNA was released into solution by alkaline digestion and subsequently precipitated. Prior to this step, the streptavidin beads were washed with immobilization buffer.

The alkaline digest was performed by addition of 100 μL 0.15 M NaOH to the beads and incubation at 55 °C for 25 min. The columns were centrifuged at 17,000 x *g*, rt for 1 min. The flow-throughs were collected in 1.5 mL DNA LoBind tubes (Eppendorf). The beads were washed by addition of 100 μL RNase-free water and centrifugation at 17,000 x *g*, rt for 1 min. Both flow-throughs were combined and neutralized by addition of 25 μL 3 M sodium acetate.

The cDNAs were precipitated in the presence of 40 μg molecular-grade glycogen and 500 μL ice-cold ethanol at −20 °C overnight. Following recovery by centrifugation at 21,000 × *g* and 4 °C for 2 h, the cDNA pellets were resuspended in 19 μL TdT reaction mixture (1 × TdT buffer, 1.25 mM CTP) and subjected to terminal deoxynucleotidyl transferase-mediated C-tailing to enable downstream PCR amplification.

Subsequently, 1 μL (20 U) TdT was added, and reactions were incubated at 37 °C for 30 min followed by heat inactivation at 70 °C for 10 min. The resulting C-tailed cDNAs were then subjected to ligation of a second adaptor required for downstream PCR amplification. The ligation mixture was prepared on ice in an 80 μL volume containing 1 × standard ligation buffer, 5 μM each pre-annealed cDNA anchor forward and reverse oligonucleotides, 10 μM ATP, 1.5 Weiss U/μL T4 DNA ligase HC (Thermo Fisher Scientific), and the complete 20 μL TdT reaction.

The mixtures were incubated at 4 °C overnight, were inactivated by heating to 65 °C for 10 min and were precipitated by addition of 20 μL 3 M sodium acetate, 100 μL H_2_O, 2 μL (40 μg) glycogen molecular grade and 500 μL 100 % ethanol and incubation overnight at −20 °C and subsequent centrifugation. cDNA pellets were dissolved in 30 μL RNase-free water.

To determine the optimal number of amplification cycles and avoid PCR overamplification, test PCRs were subsequently performed on the recovered cDNA. Reactions were carried out in a 50 μL volume using 1.0 μL precipitated cDNA in the presence of 1 × High-Fidelity PCR Master Mix (NEB), 2 μM forward primer with native barcode, and 2 μM reverse primer with native barcode. PCR products were analysed after 10, 12, 14, 16, 18, 20, 22 and 24 cycles by 2 % agarose gel electrophoresis.

Based on the test PCRs, 15 amplification cycles were chosen for final library generation. Accordingly, cDNA libraries were amplified in 100 µL reactions containing 2.0 µL precipitated cDNA, 1 × High-Fidelity PCR Master Mix (NEB), and 2 µM each barcoded forward and reverse primer (e.g., BC1 forward and reverse; Supplementary Table S2). PCR products were generated using 15 amplification cycles and subsequently subjected to native PAGE purification. The PCR reactions were purified by 10 % native PAGE. dsDNA was visualized using the Typhoon scanner and the smear was excised above 150 bp. dsDNA was eluted from the gel material by incubation in 4-5 mL 0.3 M NaOAc pH 5.5 while shaking at 350 rpm, 20 °C overnight. Then, the eluates were filtered through small 0.45 μm filters by recurring centrifugation at 5,600 x *g*, for 1 min. Filtered dsDNAs were precipitated by adding 2.5 x volumes of ice-cold ethanol, incubating at −20 °C and centrifuging at 18,500 x *g* at 4 °C for 2 h. Supernatants were discarded, and dsDNA-pellets were air-dried for a few minutes and resuspended in 200 μL RNase-free water. dsDNAs were ethanol-precipitated again and dsDNA pellets were resuspended in 10 μL RNase-free water. DNA concentrations were measured using the Quantus Fluorometer and QuantiFluor dsDNA System (Promega Corporation). Equimolar amounts (∼ 33 fmol) of each sample were pooled for the subsequent Nanopore sequencing. The final volume was reduced to 25 µL using a SpeedVac concentrator. Enrichment libraries were subsequently converted into Nanopore sequencing libraries using the SQK-LSK109 Ligation Sequencing Kit (Oxford Nanopore Technologies) according to the manufacturer’s guidelines, starting with 390 fmol input dsDNA.

Sequencing was performed on Flongle flow cells (R9.4.1 chemistry) on a MinION device. Sequencing data of biological duplicates comprising six infection time points are available under BioProject PRJNA1073512. To experimentally confirm differential enrichment observed by NAD captureSeq, selected candidate transcripts were analyzed by qPCR. Corresponding primer pairs were designed using the Primer3 web tool with a target annealing temperature of 59 °C (Untergasser et al. 2012). 1 µl of 1:50-diluted, ligated cDNAs (input material for final PCR before sequencing) were run in the presence of 500 nM qPCR primers and 1x iTaq Universal SYBR Green Supermix (Bio-Rad) and in technical duplicates in 20 µl scale. Therefore, the CFX384 Touch Real-Time PCR detection system (Bio-Rad) was used with an initial denaturation of 30 s at 95 °C followed by 50 cycles of denaturation at 95 °C for 5 s and elongation at 60 °C for 30 s following the recording of a denaturation curve to evaluate amplicon specificity.

### NAD captureSeq data analysis

Nanopore sequencing data was basecalled using the guppy basecaller (version 6.1.3). Reads were demultiplexed and trimmed with Porechop (version 0.2.4, https://github.com/rrwick/Porechop). Quality control was performed using pycoQC (Leger and Leonardi 2019). Full-length reads were subsequently detected and re-oriented, if the reads belonged to the reverse strand (classification) by Pychopper (version 2.5.0, https://github.com/epi2me-labs/pychopper). Classified or non-classified reads were subsequently mapped to the reference genomes of *E. coli* K12 (U00096.3, https://www.ncbi.nlm.nih.gov/nuccore/U00096.3), T4 phage (NC_00086.4, https://www.ncbi.nlm.nih.gov/nuccore/NC_000866.4) and 100 nt control RNA (spike-in, Supplementary Table S3) with minimap2 (version 2.21) (Li 2018). As in previous work (Wolfram-Schauerte et al. 2023), we used the *E. coli* K12 reference genome, since a matched, well-annotated reference for our *E. coli* B strain is lacking and mapping and feature assignment was not affected by this choice. The resulting sam files were sorted, filtered for primary alignments and converted to bam files using samtools (version 1.7) (Danecek et al. 2021). Reads mapping to features were counted using featureCounts (from subread package version 2.0.1) (Liao et al. 2014) allowing for multioverlapping reads and using the respective gff3 annotation files from NCBI as well as a self-composed annotation file for the 100 nt control RNA as reference annotations for the feature “gene”. Reads were subjected to manual inspection in Integrative Genomics Viewer (IGV, version 2.13.0) (Thorvaldsdottir et al. 2013).

Counted reads were further analyzed using R (version 4.1.2) using ggplot2 (version 3.3.5), dplyr (version 1.0.7), tidyverse (version 1.3.1), reshape (version 0.8.8) and VennDiagram (version 1.7.3). The custom script is available at https://github.com/MaikTungsten/PhageEpitranscriptomics. For analysis of promoters of identified NAD-RNAs, the following approach was used. The TSSs were identified using IGV. Briefly, a TSS was defined based on genome coverage either as the position, where coverage has steeply increased, or - in case of a modest increase of coverage - as the position where coverage for the NAD-RNA is first detected while trimming off initial C or G considered as artefacts from library preparation. Promoters were extracted from the reference genomes encompassing the TSS (+1) and 50 nucleotides upstream. Then, collected promoter sequences were used to generate specified MEME motifs including all sequences across all 51 positions in each motif (Bailey et al. 2015).

### dRNA-Seq library prep

Differential RNA-Seq (dRNA-Seq) was performed on total RNA from a single biological replicate including the time points t0, t4, t10 and t20 isolated as described above. dRNA-Seq library prep was conducted by vertis Biotechnologie AG. Briefly, RNA was fragmented by sonication, followed by treatment with polynucleotide kinase (NEB). Subsequently, half of the RNA was subjected to TEX treatment, whilst the other half was mock-treated. The RNA samples were polyA-tailed and 5’-PPP-ends were trimmed to 5’-P-ends via RNA 5’-polyphosphatase (Epicentre). First-strand cDNA synthesis was performed using an oligo(dT)-adapter and M-MLV reverse transcriptase followed by high-fidelity PCR amplification of the cDNA using primers suitable for Illumina TruSeq sequencing. Resulting dsDNA was purified with Agencourt AMPure XP beads (Beckman Coulter Genomics) and analysed by capillary gel electrophoresis (Shimadzu MultiNA microchip electrophoresis system). Equimolar amounts of each sample were pooled and size fractionated yielding a size range of 200 – 600 bp. The final library pool was sequenced on an Illumina NextSeq 500 system using 75 bp read length. dRNA-Seq data is available under accession code GSE255091 at the GEO database.

### dRNA-Seq data analysis

Data analysis was performed in a miniconda environment with Python 3.9. Data quality was confirmed using FastQC (version 0.11.9). Reads from fastq files were mapped to the reference genomes of *E. coli* K12 (U00096.3, https://www.ncbi.nlm.nih.gov/nuccore/545778205, accessed 10^th^ October 2022) and T4 phage (NC_000866.4, https://www.ncbi.nlm.nih.gov/nuccore/nc_000866.4, accessed 10^th^ October 2022), which were retrieved from NCBI database, using the READemption pipeline (version 2.0.3) at standard settings (Förstner et al. 2014). Alignments were converted to wiggle files using the coverage function of READemption (Förstner et al. 2014).

Wiggle files from READemption pipeline were used as input for transcription start site prediction with Annogesic (Docker image retrieved 15^th^ November 2022 from https://hub.docker.com/r/silasysh/annogesic/), which makes use of TSS predator (Dugar et al. 2013; Yu et al. 2018). In order to optimize the prediction parameters, transcription start site annotations of *E. coli* U00096.3 were retrieved from RegulonDB (accessed 13^th^ November 2022, https://regulondb.ccg.unam.mx/menu/download/datasets/index.jsp) and the TSSs from the first 200,000 genomic bases were gathered in a gff3 annotation file. The reference TSS file and the wiggle files for *E. coli* K12 and t0 were used to run TSS optimization with Annogesic at standard settings. The optimized prediction parameters were used to predict TSSs for all time points for T4 phage and *E. coli* wiggle files. Promoter motifs were generated as described above for NAD-RNA promoters.

### Comparison of dRNA-Seq and NAD captureSeq data

TSSs of NAD-capped transcripts called by the parameters mentioned above were retrieved from alignments in IGV and gathered in a gff3 file for T4 phage and *E. coli* separately. A custom R Script (R version 4.2.1) was compiled to search for the nearest TSS (within +/- 8 bp window), respectively, generated by TSS prediction based on the dRNA-Seq data. The custom script is available under https://github.com/MaikTungsten/PhageEpitranscriptomics.

### NAD-capping of purchased RNAs *in vitro*

Capping reactions were performed in a 50 µL scale using 40 µM 5’-P-RNA-Cy5 (Supplementary Table S3), 50 mM MgCl_2_ and approx. 5 mg of imidazolide nicotinamide mononucleotide (Im-NMN) as previously described (Höfer et al. 2016a). Reactions were purified by centrifugation at 14,000 x *g*, 4 °C in 0.5 mL 3 kDa Amicon filters and washing with four column volumes of RNase-free water. RNAs were eluted in a final volume of 50 µL and stored at – 20 °C. RNA concentrations and NAD-capping efficiencies were determined by 20 % PAGE or 20 % APB-PAGE analysis using unmodified input RNA as a reference and ImageLab 6.1 for quantification of band intensities. NAD-capping efficiencies accounted for around 50 %.

### Synthesis of ^32^P-labelled NAD

500 µM nicotinamide mononucleotide (NMN), 10 mM DTT, 1 µCi/µL alpha-^32^P-ATP (Hartmann Analytic), 1 x degradation buffer and 2.14 µM NadR (self-purified) were incubated at 37 °C for 1 h. The reaction was purified by P/C/I extraction and monitored by thin layer chromatography (TLC) using a 60:40 mixture of 100 % ethanol and 1 M NH_4_OAc, pH 5 as mobile and Alugram TLC plates (Macherey Nagel) as stationary phase.

### Construction of vectors for protein expression and protein expression

The *nudE.1* (GeneID: 1258692; UniprotID: P32271, Suplementary Table S2) was amplified from T4 phage DNA by high-fidelity PCR using forward and reverse primers introducing sites for restriction-based cloning as indicated in Supplementary Table S2. PCR products, purified via QIAQuick PCR purification kit (Qiagen), and pET28a vector, purified via GeneJet Plasmid Mini Prep Kit (Thermo Fisher), were both digested with NcoI and XhoI (both Thermo Fisher Scientific) and purified via PCR purification or gel extraction kit (both Qiagen). Vector and insert were ligated using T4 DNA ligase (Thermo Fisher Scientific) according to manufacturer’s instructions and ligation products transformed into chemically competent *E. coli* BL21 DE3 cells. Plasmids isolated from clones were validated for correct insert via Sanger sequencing (Microsynth Seqlab) and finally transformed into chemically competent *E. coli* BL21 DE3 cells again for protein expression (Supplementary Table S4).

A similar workflow was employed for site-directed mutagenesis of plasmids replacing the restriction-based approach with a primer-driven site-specific mutagenesis approach with 5’-monophosphorylated primers by PCR amplification and re-ligation of the PCR product with mutation. The plasmids containing the desired mutations were retrieved as described above. To assess the biochemical properties of the encoded proteins, recombinant protein expression and purification were subsequently performed.

Proteins were expressed from the respective plasmids (Supplementary Tables S1 and S4) in *E. coli* BL21(DE3). Cells were grown in LB medium at 37 °C, 180 rpm to an OD_600_ of 0.8, when protein expression was induced by addition of Isopropyl-beta-D-thiogalactoside to a final concentration of 1 mM. Bacteria were pelleted after incubation at 37 °C for 3 h. Pelleted bacteria were resuspended in Ni-NTA buffer A (50 mM Tris-HCl, pH 7.5, 1 M NaCl, 1 M urea, 5 mM MgSO_4_, 5 mM β-mercaptoethanol, 5 % glycerol, 5 mM imidazole, one tablet complete EDTA-free protease inhibitor cocktail (Roche) per L) and lysed by sonication (2 x 5 min at 80 % amplitude, 0.5 s pulse). The lysate was cleared by ultra-centrifugation at 37,000 x *g*, 30 min, 4 °C and the supernatant was filtered through 0.45 µm filters.

Proteins were purified from the supernatant by Ni-NTA affinity chromatography using either Ni-NTA agarose beads (Jena Bioscience, for gravity-based purification) or 1 mL Ni-NTA HisTrap column (GE HealthCare, for fast performance liquid chromatography (FPLC)-based purification). Proteins were eluted either with Ni-NTA buffer B (with 300 mM imidazole added, gravity-based) or a gradient using Ni-NTA buffer B (FPLC-based) and analysed by SDS-PAGE. An NGC system (Bio-Rad) was used for all FPLC-based protein purifications.

Proteins were further purified by size-exclusion chromatography (SEC) using a Superdex 200 10/300 GL column (GE HealthCare) integrated in the NGC system. SEC buffer containing 300 mM NaCl and 50 mM Tris-HCl, pH 7.5 was used as running buffer. Fractions of interest were analysed by SDS–PAGE, pooled and concentrated in Amicon Ultra-4 centrifugal filters (molecular weight cut-off (MWCO) 10 kDa with centrifugation at 5,000 x *g*, 4 °C). Protein concentration was measured with a NanoDrop ND-1000 spectrophotometer (Thermo Fisher Sceintific). Finally, proteins were stored in SEC buffer supplemented with 50 % glycerol at −20 °C. For the estimation of the oligomeric state of the proteins, 150 µL protein analysed on a Superdex 200 10/300 GL column (GE HealthCare) integrated in an NGC system via constant flow of SEC buffer with known monomeric proteins serving as calibration standards. Python package seaborn was used to perform linear regression of calibration standards and estimate the oligomeric state of analysed proteins.

### Structure predictions of NudE.1

The protein sequences of *nudE.1* WT (Supplementary Table S3) were used to predict structures via ColabFold (version 1.5.2, access date 20^th^ June 2023) (Mirdita et al. 2022). Standard settings were applied expecting NudE.1 to occur as a monomer. Alphafold3 modelling Prediction from rank 1 was further analysed. Structure visualizations and structural alignments were performed in PyMol (version 2.4.1).

### *In vitro* decapping and hydrolysis assays

500 nM NAD-capped RNA-Cy5 (Supplementary Table S3) were incubated in the presence of 1 mM DTT, 1 x degradation buffer and either 25 nM NudE.1 WT or NudE.1 E64,65Q or NudC WT (NEB). For NAD spike-in kinetics, NAD was included in the reaction at final concentrations of 350 µM (700-fold molar excess over NAD-RNA) or 750 µM (1,500-fold molar excess over NAD-RNA) and a spike-in of radioactive ^32^P-NAD (if indicated, 2.6 nM). Reactions were incubated at 37 °C and samples taken throughout time-course of the reaction were immediately stopped by addition of one volume 2 x APB loading dye (8.3 M urea, 0.05 % (w/v) bromophenol blue, 0.05 % (w/v) xylene cyanol). A sample was taken before addition of enzyme each. Samples were analyzed by 20 % APB-PAGE and gels were imaged at ChemiDoc (Bio-Rad) using the Cy5 channel. Reactions using ^32^P-NAD were stopped by heating samples at 90 °C for 1 min and were subsequently analysed by TLC. TLC plates were imaged by autoradiography using a Typhoon Imager (Amersham Biosciences). Intensity-based band quantification was performed in ImageLab (Bio-Rad).

NAD only hydrolysis assays were performed using the same settings with the following abberations. A final concentration of 25 µM NAD and a spike-in of radioactive ^32^P-NAD (2.6 nM) were used instead of NAD-RNA-Cy5 in the presence of 1 µM NudE.1 WT or NudE.1 E64,65Q or NudC WT. Reactions were analysed by TLC and autoradiography as described above.

### Determination of cellular metabolite/cofactor levels

Cultures were quenched by adding 1 mL culture to 1 mL 70 % methanol at −80 °C and harvested by centrifugation at 13,000 x *g*, −5 °C, 5 min. Supernatant was carefully removed and pellets stored at −80 °C. Intracellular metabolites were extracted adding a mixture of 50 % methanol and 50 % TE buffer pH 7 (10 mM Trizma base and 1 mM EDTA) at −20 °C to the frozen cell pellets. The volume of extraction fluid per cell was kept constant throughout all experiment by adjusting it according to the amount of cells in each pellet (volume per pellet calculated according to OD_600_ measurement). For membrane desintegration, an equal volume of −20 °C chloroform was added to each pellet, pellets were resuspended by vortexing, and extraction was performed by shaking at 4 °C, 1,000 rpm for 2 h. Organic and aqueous phase were separated by centrifugation at 21,000 x *g*, −10 °C for 10 min and aqueous upper phase was filtered through 0.2 µm PFTE filters (Phenomenex) and stored at −80 °C until analysis.

Quantitative determination of NAD, FAD, and UDP-GlcNac was performed using a LC-MS/MS. Chromatography was performed on an Agilent Infinity II 1290 HPLC system using a SeQuant ZIC-pHILIC column (150 × 2.1 mm, 5 μm particle size, peek coated, Merck) connected to a guard column of similar specificity (20 × 2.1 mm, 5 μm particle size, Phenomenex) at a constant flow rate of 0.1 mL/min with mobile phase composed of fractions of buffer A (10 mM ammonium acetate in water, pH 9, supplemented with 5 µM medronic acid) and buffer B (10 mM ammonium acetate in 90:10 acetonitrile to water, pH 9, supplemented with 5 µM medronic acid) at 40 °C.

2 µL sample were injected. The mobile phase profile consisted of the following steps of linear gradients: 0 – 1 min constant at 75 % B; 1 – 6 min linear gradient from 75 to 40 % B; 6 to 9 min constant at 40 % B; 9 – 9.1 min linear gradient from 40 to 75 % B; 9.1 to 20 min constant at 75 % B. An Agilent 6495 ion funnel mass spectrometer was used in negative and positive ionisation mode with an electrospray ionization source and the following conditions: ESI spray voltage 3500 V, nozzle voltage 1000 V, sheath gas 300 °C at 9 L/min, nebulizer pressure 20 psig and drying gas 100 °C at 11 L/min. Compounds were identified based on their mass transition and retention time compared to standards (Supplementary Table 5). Chromatograms were integrated using MassHunter software (Agilent Technologies). Absolute concentrations were determined based on an external standard curve.

According to our Coulter counter measurement an average *E. coli* culture at an OD_600_ of 0.8 contains 3 * 10^8^ cells per 1 mL (cells_per extraction volume_). Applying an average single cell volume (V_cell_) of 1 fL per cell (Kubitschek 1990; Volkmer and Heinemann 2011), the intracellular volume of all cells in a pellet derived from 1 mL of sample at an OD of 0.8 thus amounts to 3 * 10^8^ fL (=3 * 10^-7^ L). As all metabolites detected in the extract are derived from that volume, the intracellular concentration can be calculated by determining the number of metabolites in the extract (V_extract_ * c_metabolite_) using a correction factor of 10^-12^ to yield a molar concentration and divided by the intracellular volume of all cells in the pellet (prior to extraction). Thus, the concentration of each metabolite per cell (c_per cell_ [M]) can be calculated with equation (2):

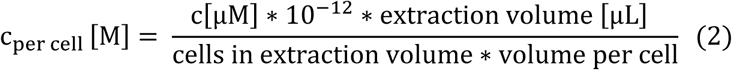

### CircNC-like protocol for NAD-RNA validation

The circNC-like protocol was conducted similar as described by Sharma and colleagues (Sharma et al. 2023). In a 100 µL scale, 8 µg total RNA from *E. coli* JM109 + pUC19 or *E. coli* B strain was dephosphorylated with 0.4 U/µL QuickCIP (NEB) in 1 x rCutSmart buffer (NEB) at 37 °C for 1 h. The RNA was purified using the Zymo RNA Clean and Concentrator kit eluting the RNA in 15 µL. Subsequently, 4 µL dephosphorylated RNA were subjected to NudC treatment by incubation in the presence of 1 mM DTT, 1 x degradation buffer (see composition above) and 1.5 µM NudC (or an equivalent volume of RNase-free water as negative control) in 50 µL scale at 37 °C for 1 h. A non-enzyme sample serves as negative control. RNA was then purified again and the eluate was subjected to circularization with T4 RNA Ligase 1 (NEB). Briefly, 15 µL RNA was incubated in the presence of 10 % PEG8000, 0.5 U/µL T4 RNA Ligase 1, 1 U/µL murine RNase inhibitor (NEB), 50 µM ATP and 1 x RNA ligation buffer (NEB) at 25 °C for 16 h. RNA was purified again and concentration determined at NanoDrop. 50 ng purified RNA was then subjected to reverse transcription with Superscript IV (Invitrogen) according to manufacturer’s instructions using 100 nM RNA-specific reverse transcription primer, followed by RNase H digest with 5 µL RNase H (NEB) at 37 °C for 20 min. RT reaction were used as template for qPCR in 10 µL scale including 300 nM forward and reverse primers and 1 x iTaq Universal SYBR Green Supermix (Bio-Rad). Using the CFX384 Touch Real-Time PCR detection system (Bio-Rad) with an initial denaturation of 1 min at 95 °C followed by 50 cycles of denaturation at 95 °C for 10 s and elongation at 59 °C for 30 s following the recording of a denaturation curve to evaluate amplicon specificity.

Log2 Fold Changes were calculated as the difference between the cq-value of the control sample and the NudC WT treated sample. cDNAs were diluted when necessary if Cq values were lower than 15, as values below this threshold are generally considered outside the reliable dynamic range for quantitative PCR. Each target was measured in technical duplicates and biological triplicates, and the mean values were calculated. Targets were quantified individually.

### Comparative Quantification of 5’-Modification Status of Selected Transcripts via CapPro Assay

To determine the 5’-phosphorylation state and modification status (PPP/NAD) of selected transcripts, total RNA was isolated from *E. coli* JM109 + pUC19 or *E. coli* B strain at 0, 10 and 20 minutes post-infection. For each sample, 50 µg of total RNA was used as input, supplemented with a spike-in of NAD-RNA (100 nt-control Qbeta-RNA) as a control.

For direct quantification of 5’-monophosphorylated RNA (P-RNA), 8 µg of total RNA were processed using the circNC-like assay, as described above. In this approach, P-RNAs were directly circularized via ligation, followed by reverse transcription and qPCR to quantify the target RNA. To distinguish 5’-triphosphorylated RNA (PPP-RNA) from other RNA species, the remaining total RNA was subjected to a sequential enzymatic treatment workflow. First, the One ScriptCap m^7^G Capping System (Cellscript) was used to cap all PPP-RNAs with an m^7^G cap. Reactions were prepared according to the manufacturers protocol and incubated at 37 °C for 2 h, followed by overnight RNA precipitation. The precipitated RNA was then treated with calf intestinal phosphatase (CIP, NEB) to dephosphorylate 5’-diphosphorylated RNAs, while the m^7^G-capped RNAs remained unchanged. For this, the samples were incubated at 37 °C for 1 h with 1 µL CIP per 1 µg RNA in the presence of 1 x Cut Smart buffer, following the manufacturers protocol. The treated RNA was then purified using Zymo RCC-5 columns and the cleaned RNA was eluted in 25 µL RNase-free water.

For direct quantification of 5’-diphosphorylated RNA (PP-RNA), 8 µg of total RNA were processed using the circNC-like assay, as described above. In this approach, P-RNAs were directly circularized via ligation, followed by reverse transcription and qPCR to quantify the initial 5’-diphosphorylated state of the target RNA.

Subsequently, the RNA was incubated with yDcpS (NEB), a decapping enzyme that specifically hydrolyzes m^7^G-capped RNAs, generating dephosphorylated 5′ -ends. Since the capacity of yDcpS is limited to 1–5 µg RNA per reaction, samples were split into multiple reactions, each containing 3 µg RNA. Incubations were performed with the reaction buffer provided by the manufacturer, following the manufacturer’s protocol, for 1 h at 37 °C. The enzyme was then inactivated by heating the reactions at 70 °C for 15 min. Immediately afterwards, RNA 5′ polyphosphatase (Biozym) treatment was carried out to convert the remaining phosphorylated RNA species into monophosphorylated RNA. The same split-reaction setup used for yDcpS treatment was applied. Samples were incubated with 1 × reaction buffer and RNA 5′ polyphosphatase for 30 min at 37 °C, according to the manufacturer’s instructions. Following enzyme treatment, RNA samples were precipitated overnight. At this stage, the initially triphosphorylated RNAs in the sample were converted into monophosphorylated RNAs, allowing their quantification via the circNC assay in combination with RT-qPCR by using 8 µg of processed RNA samples.

To quantify NAD-RNAs, an additional fraction of the remaining total RNA was treated with the alkaline phosphatase QuickCIP (NEB). Samples were incubated with QuickCIP at 37 °C for 1 h following the manufacturer’s protocol. The RNA was then purified using Zymo RCC-5 columns according to the manufacturer’s instructions, and eluted in 25 µL RNase-free water. This step is critical to ensure complete removal of phosphorylated RNAs, which could otherwise interfere with NAD-capped RNA analysis. To quantify NAD-RNAs, the remaining total RNA was treated with NudC, an NAD-decapping enzyme. For NudC treatment, reactions (50 µL total volume) contained 1 mM DTT, 1× degradation buffer (see composition above), and 1.5 µM NudC were incubated at 37 °C for 1 h. The resulting RNA was subsequently subjected to ligation using the circNC assay and analyzed by RT-qPCR to determine the initial amount of NAD-RNA present in the sample.

### Quantification of NAD-RNA using the FluorCapQ Assay

Quantification of NAD-modified RNA was performed using a fluorescence-based assay, previously published as FluorCapQ Assay (Wiedermannova et al. 2024). Total RNA was isolated from *E. coli* B strain infected with either T4 phage WT or T4 NudE.1 E64,65Q mutant at time points 0, 7 and 20 minutes post-infection. To eliminate interference from free NAD, total RNA samples as well as *in vitro* transcribed (IVT) NAD-RNA and PPP-RNA were purified via Urea treatment and filtration using 3 kDa Amicon Ultra centrifugal filters. The 3 kDa filters were equilibrated by repeated washing steps with 0.15 M NaOH and Milli-Q (MQ) water through centrifugation at 14,000 × *g* for 10 minutes at 18 °C. Filters were stored at 4 °C until use. For each sample, RNA was incubated with 8.3 M urea for 10 minutes at room temperature, followed by loading onto the equilibrated 0.5 mL 3 kDa Amicon filter. After each centrifugation step, flowthrough was discarded and new buffer added. RNA was retained in the upper chamber, and gentle pipetting ensured mixing without disrupting the membrane. Washing steps were performed as follow: 2 x with 400 µL 8.3 M Urea, 1 x with 400 µL MQ water, 2 x with 400 µL 4.15 M Urea and 4 x with 400 µL MQ water. Finally, RNA was recovered by two additional washes with 200 µL sterile water. The RNA solution (approx. 400 µL) was concentrated via vacuum centrifugation. RNA concentration was determined using Nanodrop, and RNA integrity was validated using PAGE analysis.

For the FluorCapQ assay, 10 µg of RNA was dissolved in 40 µL RNase-free water, followed by the addition of 15 µL 2-acetylbenzofuran (25 mM in ethanol) and 15 µL of 0.5 M potassium hydroxide (KOH). The reaction was incubated on ice for 20 minutes, followed by the addition of 70 µL formic acid. Samples were then incubated at room temperature for an additional 20 minutes. The resulting fluorophore was stable in the dark at room temperature for up to 2 hours. For the calibration, a standard curve was prepared in parallel using a dilution series of NAD (500 fmol to 0.5 fmol) under the same reaction conditions. All samples and standards were transferred to a black 96-well microplate, and fluorescence was measured using a plate reader (Agilent Technologies BioTek Synergy H1 multimode reader), with excitation at 420 ± 40 nm and emission at 480 ± 40 nm.

### Data visualization and statistics

If not indicated otherwise, statistical tests and plots were performed with R (version 4.2.2; R Core Team 2022) using ggplot2 (Wickham, 2016) (version 3.4.1) and ggpubr (Kassambara, 2023) (version 0.6.0). Fits of enzyme kinetics were performed in R using a generalized linear model. Promoter analyses were performed in python (version 3.11.4) using pandas (version 1.5.3; McKinney, 2010) and numpy (version 1.25.0; Harris et al., 2020) and motifs generated using meme command line tool as described above (version 5.5.3; Bailey et al., 2015).

## RESULTS

### T4 infection reduces cellular NAD and remodels NAD-RNA abundance

Since NAD is both a metabolite and an RNA-cap on bacterial RNA, we first asked whether T4 phage infection of *E. coli* changes cellular NAD pools and global NAD-RNA abundance. For this, we first assessed potential changes in the metabolome of T4 phage-infected *E. coli*. We measured the endometabolome of T4 infected *E. coli* using LC-MS. We focused at NAD and two other metabolites that serve as bacterial cofactor caps, namely FAD and UDP-N-acetylglucosamine (UDP-GlcNAc) (Julius and Yuzenkova 2017; Wang et al. 2019; Schauerte et al. 2021), to monitor potential influences of T4 phage infection on the various cofactor-caps existing in a bacterial cell. We quantified all three metabolites per *E. coli* cell before (t0) and at two time points post infection (t10, t20) (Figure 1A). On average, we detected 1.01 x 10^-4^ M FAD, 2.34 x 10^-3^ M NAD and 1.09 x 10^-3^ M UDP-GlcNAc per *E. coli* cell before infection (t0). The measured FAD and NAD concentrations agree well with previously reported metabolite concentrations in *E. coli* (FAD: 1.7 x 10^-4^ M; NAD: 2.6 x 10^-3^ M) (Bennett et al. 2009). UDP-GlcNAc concentrations deviate from previously reported values by approx. 9-fold (UDP-GlcNAc: 9.6 x 10^-3^ M) (Bennett et al. 2009). Upon T4 phage infection the FAD pool did not seem to be affected compared to uninfected *E. coli* (Supplementary Figure S1A). In contrast, we detected a slight increase of UDP-GlcNAc levels over the time course of infection (Supplementary Figure S1B). Interestingly, the NAD pool significantly decreased by almost half upon T4 phage infection (still at approx. 1.5 mM) while remaining constant in uninfected *E. coli* (Figure 1A). In comparison, several phage defense systems employ NAD depletion to abort infection (Ofir et al. 2021; Garb et al. 2022; Zaremba et al. 2022), which reduce NAD levels by almost half to 5-fold (Ofir et al. 2021; Garb et al. 2022). Thus, the NAD levels in *E. coli* decrease in a similar manner during T4 phage infection as induced by some anti-phage defense systems (Osterman et al. 2024b).

**Figure 1:**
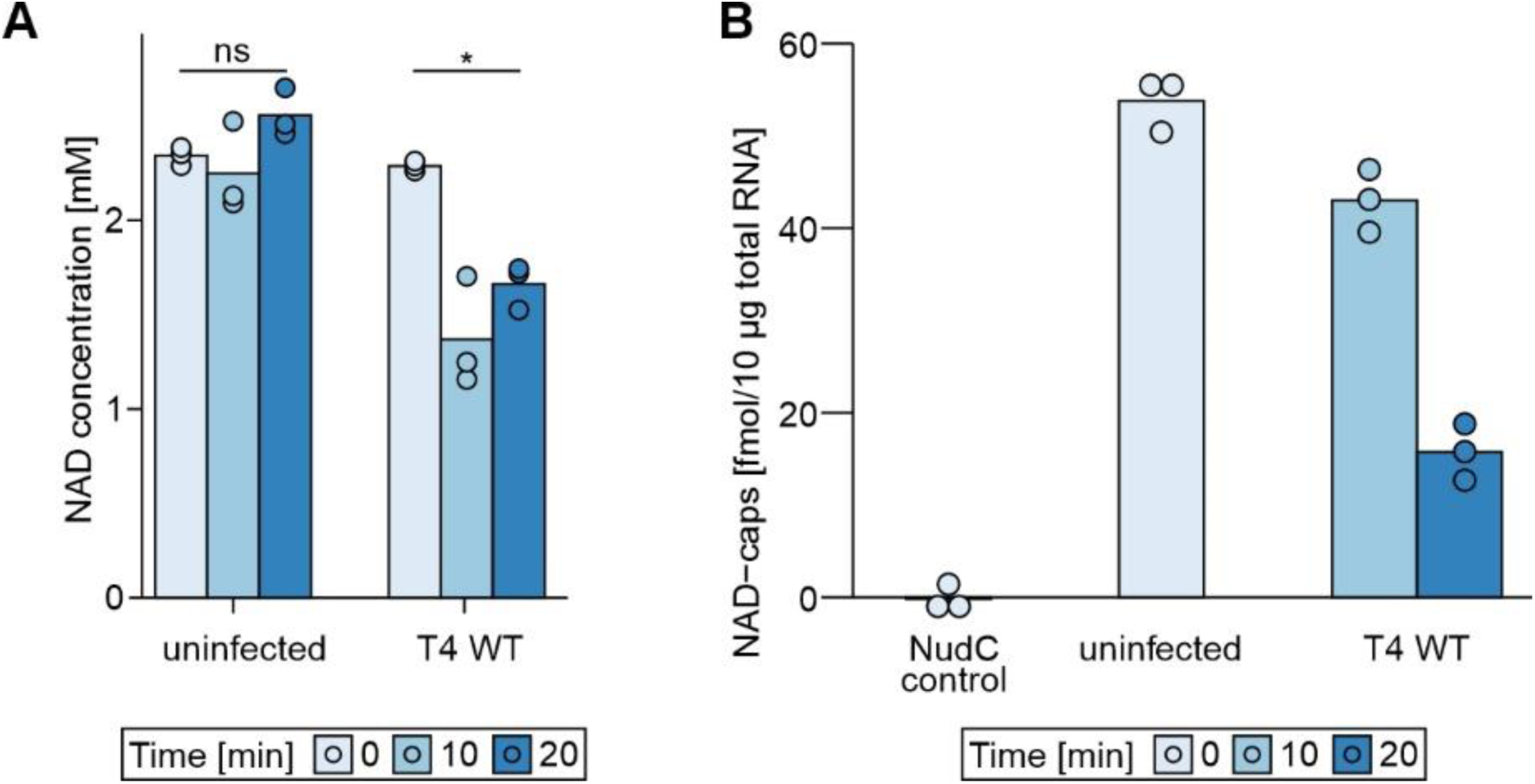
Coordinated depletion of NAD and NAD-capped RNAs during T4 phage infection. **(A)** Intracellular NAD concentrations measured by metabolomics in uninfected *E. coli* and during T4 WT infection at the indicated time points 10 and 20 minutes post infection (n = 3). NAD concentrations [mM] were calculated assuming an *E. coli* cell volume of 1 fL. A two-sided t-test comparing NAD levels at 0 and 20 minutes found changes upon T4 WT infection to be significant (*: p < 0.05), while changes in uninfected *E. coli* were non-significant (ns). **(B)** Total NAD-cap abundance determined from total RNA isolated under the same conditions. The NudC control represents total RNA treated with recombinant NudC prior to analysis and serves as a negative control, as removal of NAD caps is expected to abolish the fluorescence signal. Uninfected samples correspond to total RNA isolated from *E. coli* cells, whereas T4 WT samples correspond to total RNA isolated from *E. coli* 10 min and 20 min after infection with bacteriophage T4. T4 infection resulted in a decrease in both intracellular NAD levels and total NAD-capped RNA abundance compared to uninfected cells. Bars represent mean values and dots indicate individual biological replicates (*n* = 3). Error bars represent standard deviation (s.d.).

To assess the overall presence of NAD-capped RNAs in *E. coli* during T4 phage infection, we employed a fluorescence-based assay previously published as the FluorCapQ assay (Wiedermannova et al. 2024). Using this assay, we quantified NAD-RNA levels from total RNA isolated at 0, 7 and 20 minutes post-infection to capture the temporal dynamics of phage infection (Figure 1B). This allowed us to assess potential differences in NAD-RNA levels across uninfected *E. coli* and T4 phage infection phases. Our analysis demonstrated a significant decrease in NAD-RNA levels during T4 phage infection, with a clear reduction from approx. 54 fmol prior infection to approx. 16 fmol at 20 minutes post-infection (p < 0.001; two-sided Welch two sample t-test) (Figure 1B).

Together, these findings demonstrate that T4 phage infection induces a strong decrease of both cellular NAD levels and the NAD-RNA pool. Yet, we cannot distinguish whether only host or phage NAD-RNAs specifically or both decrease in abundance per cell, since we quantify the overall number of NAD-caps per cell.

### The NAD-RNA pool is dynamically reshaped during T4 phage infection

During T4 infection, the host transcriptome is rapidly remodeled while phage gene expression proceeds in a highly dynamic and temporally ordered manner (Wolfram-Schauerte et al. 2022) Observing the decrease in cellular NAD and NAD-cap levels, we wondered whether similar concepts may apply to the NAD-RNA pool during T4 phage infection. How are host NAD-RNAs affected by infection? Are phage transcripts NAD-capped and is this capping similarly dynamic as the expression of phage genes during infection?

In order to comprehensively determine the occurrence of both NAD-capped host and phage transcripts during infection, we made use of the NAD captureSeq technology (Cahova et al. 2015; Winz et al. 2017) adapted to sequence full-length transcripts with the Oxford Nanopore (Supplementary Figure S2). We included a 100 nt control NAD-RNA (Supplementary Table S3) from the beginning of the NAD captureSeq workflow to monitor the success of the extensive capture and sequencing protocol.

We applied our NAD captureSeq pipeline to comprehensively determine the occurrence of NAD-RNAs before (t0) and during T4 phage infection (1, 4, 7, 10, 20 min post infection) (Supplementary Figure S3). In total, we sequenced 12 samples per replicate, comprising all six time points in the presence and absence of ADPRC, and obtained 220,028 pass reads in replicate 1 and 499,383 pass reads in replicate 2 (Supplementary Figure S4). After demultiplexing, this corresponded to an average of 14,777 and 36,255 reads per sample, respectively.

We measured enrichment of the 100 nt control NAD-RNA in both replicates and all time points (Figure 2A, B, Supplementary Table S6 A, B, Supplementary Figure S5, S6) indicating that our NAD captureSeq workflow successfully captured NAD-RNAs in all samples. It is important to note that NAD-RNAs are detected based on their enrichment, and their measured abundance does not necessarily reflect their overall expression levels. Furthermore, due to the enrichment-based nature of the method, NAD captureSeq does not provide information about the relative proportion of NAD-capped transcripts compared to uncapped transcripts such as those with 5’ PPP-, 5’ P- or 5’ OH.

**Figure 2:**
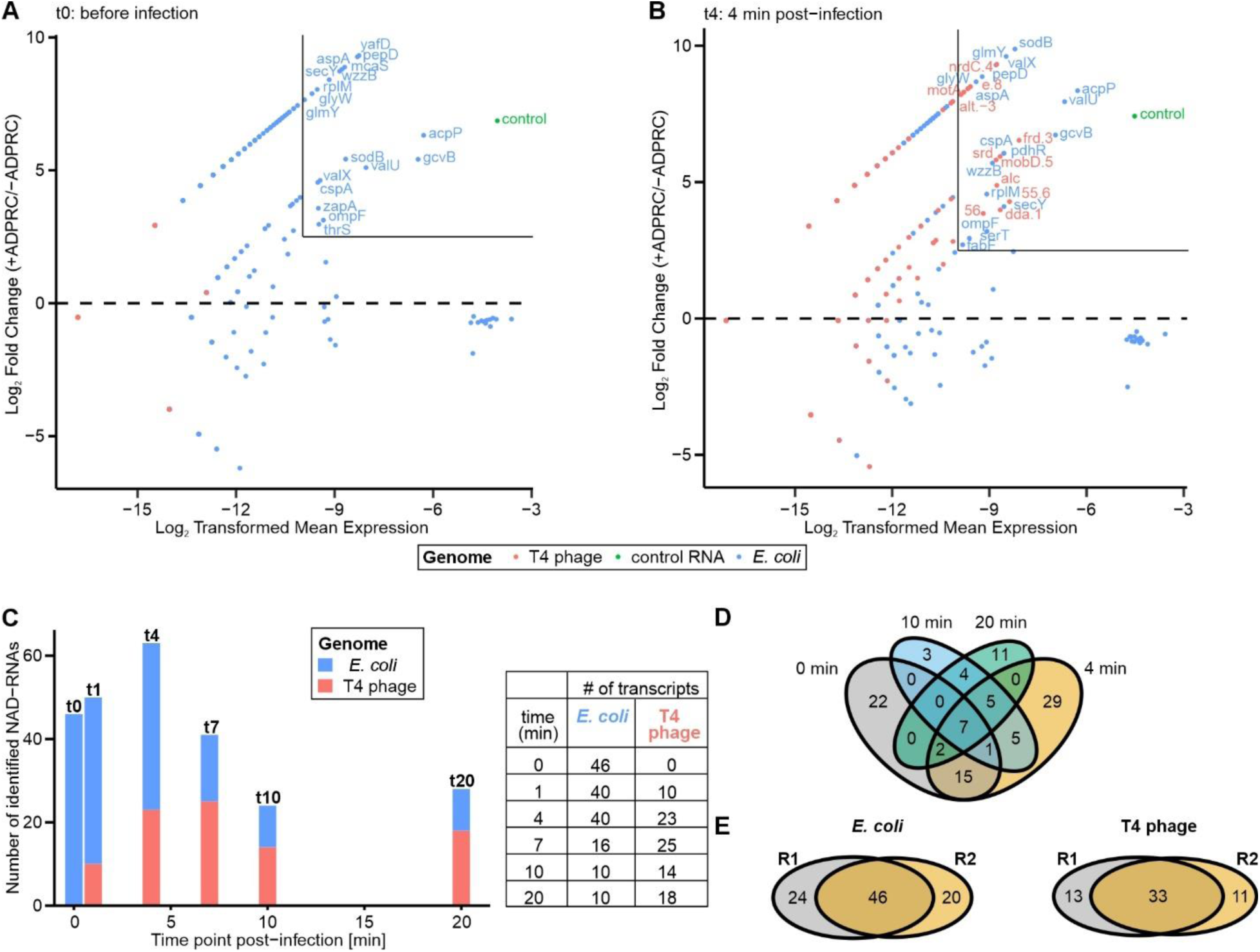
Identification of NAD-RNAs using time-resolved dual NAD captureSeq. (**A**, **B**) M (log-ratio) A (log average) plots showing enrichments of NAD-capped RNAs before infection (t0, **A**) and 4 min post infection (t4, **B**) for replicate 1 at an MOI of 1.5 (Supplementary Figure S3B). y-axis represents the log2 fold change in normalized read counts comparing fully-treated sample (+ADPRC) and negative control (-ADPRC), x-axis shows log2 transformed mean normalized read counts for genes from +ADPRC and –ADPRC samples. Plots for all time points and replicates are shown in Supplementary Figures S5 and S6. (**C**) Bar plot showing the number of NAD-capped transcripts identified by time-resolved NAD captureSeq throughout the course of T4 infection for biological replicate 1. The table on the right displays the number of *E. coli* transcripts (blue) identified as NAD-capped at 0, 1, 4, 7, 10, and 20 min post infection, and the corresponding number of NAD-capped T4 phage transcripts (peach) detected at the same time points. (**D**) Distribution of NAD-RNAs across different infection phases for replicate 1. (**E**) Distribution of *E. coli* and T4 phage NAD-RNAs identified in different replicates.

Before infection, the identified *E. coli* NAD-RNAs were consistent with previously reported NAD-capped transcripts, including regulatory RNAs such as GcvB, McaS, GlmY, and GadY, as well as mRNAs such as *acpP* and *aspA* (Cahova et al. 2015; Zhang et al. 2021). Further, we detected several *E. coli* tRNA transcripts from the *valU-valX-valY* and *metZ-metW-metV* operons as NAD-capped (Figure 2A, Supplementary Table S6A, B, Supplementary Figure S5, S6). Zhang and colleagues also identified NAD-capped tRNAs (Zhang et al. 2021), which is generally unexpected due to the post-transcriptional processing of tRNAs from an RNA precursor into mature tRNA molecules (Shepherd and Ibba 2015). Closer inspection of the read distribution at the respective tRNA loci demonstrated that tRNAs are NAD-capped as polycistronic precursor transcripts spanning the entire operon (Supplementary Figure S7). This exemplifies the power of the SPAAC- and ONT-based NAD captureSeq approach to enrich full-length NAD-capped transcripts or longer fragments thereof.

Next, we investigated how the NAD-RNA landscape changes during T4 phage infection. Consistent with previous dual-transcriptome studies showing host RNA depletion and temporally regulated phage gene expression (Luke et al. 2002; Wolfram-Schauerte et al. 2022), the number and identity of NAD-RNAs changed markedly over the course of infection.

First, the number of identified *E. coli* NAD-RNAs decreases over the time course of infection (Figure 2C) ranging from 46 and 43 RNAs before infection in replicate 1 and 2, respectively, to only 10 and 14 RNAs at 20 min post infection. In addition, we identified a variety of T4 phage transcripts as NAD-capped providing the first evidence for NAD-capped phage RNA. Throughout infection, T4 phage NAD-capped RNAs vary in their number and identity (Figure 2C). Predominantly, T4 phage mRNAs appear to be NAD-capped. The functions of NAD-RNAs vary with the respective infection phase. Early NAD-capped RNAs play roles in host takeover, whilst middle and late T4 NAD-RNAs are encoded by genes for DNA replication or structural phage proteins, respectively (Supplementary Table S6 A, B). This demonstrates the dynamic nature of the NAD-RNA landscape during T4 phage infection.

Comparing the NAD-RNAs detected over the time course of infection, we found that most *E. coli* NAD-RNAs are present before or early during infection (Figure 2D), while a small set of host NAD-RNAs could be detected across all infection phases and replicates (Table 2). In contrast, T4 NAD-RNAs are only shared by two or three adjacent time points of infection or are unique for one time point (Figure 2D, Supplementary Table S 6A, B) underlining the dynamic expression of phage NAD-RNAs.

**Table 2:** Abundant *E. coli* and T4 phage NAD-RNAs identified by NAD captureSeq. Transcripts are ranked from top to bottom according to their relative abundance, with the most enriched NAD-RNAs listed at the top.

| Abundant *E. coli* NAD-RNAs during infection |  |  |
| --- | --- | --- |
| Gene | RNA type, occurrence | Function |
| *gcvB* | sRNA, entire infection | regulation of amino acid metabolism (Urbanowski et al. 2000) |
| *glmY* | sRNA, entire infection | response to cell envelope stress (Hobbs et al. 2010); regulation of GlmZ (Reichenbach et al. 2008); stable during T4 phage infection (Wolfram-Schauerte et al. 2022) |
| *acpP* | mRNA, entire infection | acyl carrier protein in fatty acid biosynthesis (Rawlings and Cronan 1992) |
| *sodB* | mRNA, entire infection | protection from oxidative stress (Carlioz and Touati 1986) |
| *valU* | valine-tRNA, entire infection | valine-tRNA |
| **Most abundant T4 phage NAD-RNAs** |  |  |
| **Gene** | **RNA type, occurrence** | **Function** |
| *dda.1* | mRNA, middle to late phase | homing endonuclease (predicted by Phold) |
| *srd* | mRNA, middle to late phase | inducer of RNase E-mediated RNA decay (Qi et al. 2015) |
| 56 | mRNA, early to middle phase | involved in a protein complex for nucleotide dephosphorylation (Munro and Wiberg 1968) |
| 55.6 | mRNA, early to middle phase | uncharacterized protein |
| *frd.3* | mRNA, early phase | uncharacterized protein |

To further determine the robustness of the NAD captureSeq results, we compared the *E. coli* and T4 phage NAD-RNAs that were overall identified in both replicates. We observed substantial overlap between biological replicates (Figure 2E), supporting the reproducible detection of many the generation of NAD-capped host and phage transcripts. Some differences are expected as a) NAD captureSeq is not a quantitative approach relying on enrichment and b) our selected threshold for calling NAD-RNAs is set to stringently exclude low abundant NAD-RNAs to increase the confidence of detected NAD-RNAs. Due to variations in enrichment scores, NAD-RNAs may show enrichments closely above the threshold in one and below in the other replicate causing inconsistencies across replicates. Finally, we validated our data by performing qPCR on the cDNAs used for final library amplification and sequencing to exclude potential amplification or sequencing biases. We recorded enrichments for NAD-RNAs in the +ADRPC-sample compared to the -ADPRC-control as reflected in the NAD captureSeq data (Supplementary Table S7).

### Specific host and phage transcripts carry NAD-caps

We next set out to inspect specific host and phage NAD-RNAs in more detail. As described above, we found NAD-capped polycistronic host tRNA precursor transcripts consistent with previous studies (Zhang et al. 2020). The *valU*-tRNA precursor RNA is among the five host NAD-RNAs detectable over the entire time course of infection (Table 2). Two sRNAs, GcvB and GlmY, are among the host NAD-RNAs abundant during the entire infection, which are well-established NAD-capped RNAs (Cahova et al. 2015). Interestingly, GlmY is a transcript known to be expressed at rather constant abundance during T4 phage infection (Wolfram-Schauerte et al. 2022).

To provide evidence for NAD-capping of these transcripts independent from our NAD captureSeq experiment, we first employed a ligation-based assay similar to the circNC approach to independently verify selected NAD-RNAs (Sharma et al. 2023) (Supplementary Figure S8A). We validated this protocol using NAD-RNAI (Cahova et al. 2015) as positive and 5S rRNA and 6S RNA as negative controls (Cahova et al. 2015; Zhang et al. 2021) (Figure 3A). Then, we confirmed GcvB as an NAD-capped RNA in *E. coli* (Figure 3A).

**Figure 3:**
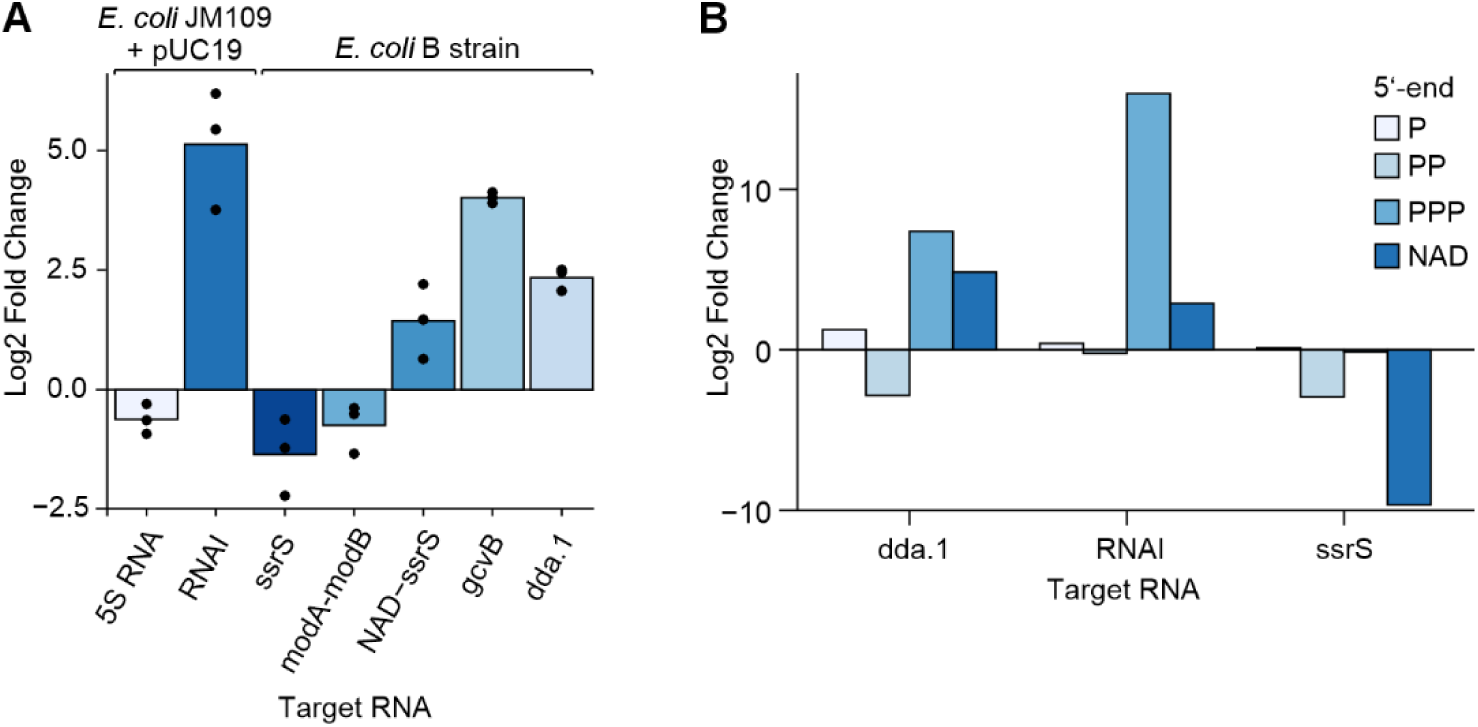
Validation of NAD-RNAs via a circNC-like and CapPro assay. (**A)** Log2 fold changes between control and NudC-treated total RNAs (*E. coli* JM109 + pUC19) in qPCR targeting selected *E. coli* and T4 phage NAD-RNAs (*E. coli* B strain) using our ligation-based assay (n=3). (**B)** Relative abundance of 5′-monophosphorylated (P), 5′-diphosphorylated (PP), 5′-triphosphorylated (PPP), and NAD-capped transcript species determined using the CapPro assay (workflow shown in Supplementary Figure S6). Transcript-specific enrichment was quantified by circNC-based circularization followed by RT-qPCR and is shown as log2 fold change for 6S RNA, RNAI, and dda.1 at the indicated time points. To validate the workflow, a synthetic 100-nt NAD-capped Qβ RNA was added as an internal spike-in control prior to RNA processing. In addition, RNAI served as a positive control, as a fraction of RNAI is known to carry an NAD cap, whereas 6S RNA served as a negative control for NAD capping. Consistent with these expectations, NAD-specific signal was detected for RNAI but not for 6S RNA, supporting the specificity of the assay and the correct discrimination of 5′-end modification states. Data are derived from a single biological replicate with two technical qPCR replicates (n = 1, technical duplicates).

Unexpectedly, NAD captureSeq also indicated specific *E. coli* NAD-RNAs, present only during but not before phage infection potentially suggesting that phage infection may induce condition-specific NAD-capped host transcripts. For instance, we found an NAD-RNA to be transcribed from an alternative TSS in the 6S RNA locus (*ssrS* gene) particularly in middle and late infection phase (7 – 20 min post infection) (Supplementary Table S6A, B). The TSS is +1A and located 60 bp downstream of the canonical TSS of the *ssrS* gene. Using the circNC-like assay, we confirmed the NAD-cap on the specific transcript, while the full-length 6S RNA is not NAD-capped (Figure 3A). This exemplifies that even though phage infection is conceived to primarily shut-down host transcription, specific host genes could be transcribed as modified transcripts.

In addition to host NAD-RNAs, we discovered the first T4 phage NAD-RNAs, among which the T4 protein-coding genes *dda.1*, *srd*, *56*, *55.6* and *frd.3* encode the most abundant T4 phage NAD-RNAs (based on detection in both replicates and highest mean expression) (Table 2). Most of the proteins encoded by these NAD-RNAs are functionally uncharacterized except for Srd that induces RNase E-mediated RNA decay (Qi et al. 2015) and 56 that is part of a complex for nucleotide dephosphorylation (Munro and Wiberg 1968). Importantly, these likely abundant T4 phage NAD-RNAs do not belong to the top 10 most highly expressed T4 genes at any time point of infection as indicated by a previous RNA-Seq study (Wolfram-Schauerte et al. 2022). The NAD-capped *dda.1* transcript showed the overall highest abundance in the NAD captureSeq experiment. We confirmed this transcript as NAD-capped with the circNC-like assay, while an uncapped transcript derived from the *modA-modB* locus was not detected as NAD-capped (Figure 3A).

The likely high abundance of the *dda.1* NAD-RNA prompted us to quantify its degree of NAD-capping. Since the circNC-like assay does not provide a quantitative measure for the fraction of NAD-capped transcript, we set out to develop the CapProAssay to systematically quantify the abundance of 5’-P, -PP, -PPP and -NAD ends for a transcript of interest (Supplementary Figure S8B). The CapProAssay enables quantitative profiling of transcript 5′ -end chemistry, including distinct phosphorylation states and NAD-caps in a unique way. For our positive control *RNAI*, we demonstrated that approx. 15 % of the transcript population was NAD-capped, which agrees with previously published data reporting 10 – 50 % NAD capping in total RNA (Cahova et al. 2015). In addition, no NAD-capping was detected for the negative control, 6S RNA, thereby confirming the specificity of our assay. Using the CapProAssay we could identify that approx. 35 % of the T4 phage *dda.1* transcript is NAD-capped, in comparison to its 5′-PPP and 5′-P counterparts (Figure 3B). Closer inspection of the *dda.1* locus revealed that it is mainly expressed in the middle phase of infection alongside three downstream genes (*dda*, *dexA.2*, *dexA.1*) (Wolfram-Schauerte et al. 2022), for which only Dda is known as a helicase involved in nucleic acid metabolism. Consequently, NAD-capping could occur on a polycistronic transcript ranging from *dda.1* to *dexA.1*. This observation prompted us to systematically investigate the mechanism of NAD-capping during T4 phage infection.

### The host RNAP installs NAD-caps on phage and host transcripts

The *E. coli* RNAP, which caps *E. coli* NAD-RNAs *ab initio* during transcription (Bird et al. 2016), is also responsible for the transcription of T4 phage genes during infection (Miller et al. 2003). Thus, paired with our observation of early, middle and late T4 NAD-RNAs (Figure 2), we supposed that T4 phage transcripts are analogously NAD-capped by the host RNAP. Therefore, we performed differential RNA-Seq (dRNA-Seq) (Sharma et al. 2010; Sharma and Vogel 2014; Wicke et al. 2021) to determine the global transcription start sites (TSSs) in *E. coli* before and at three time points of T4 phage infection. Thereby, we identified in total 5341 *E. coli* TSSs and 150 T4 phage TSSs including 22 phage antisense TSSs (Figure 4 A, B, Supplementary Table S9 A, B). The number of identified TSSs decreases for *E. coli* over the time course of infection (from 4830 TSSs (t0) to 2996 TSSs (t20)), whilst the number of T4 phage TSSs is increasing with no meaningful TSS identified prior to infection (Supplementary Table S9 A, B).

**Figure 4:**
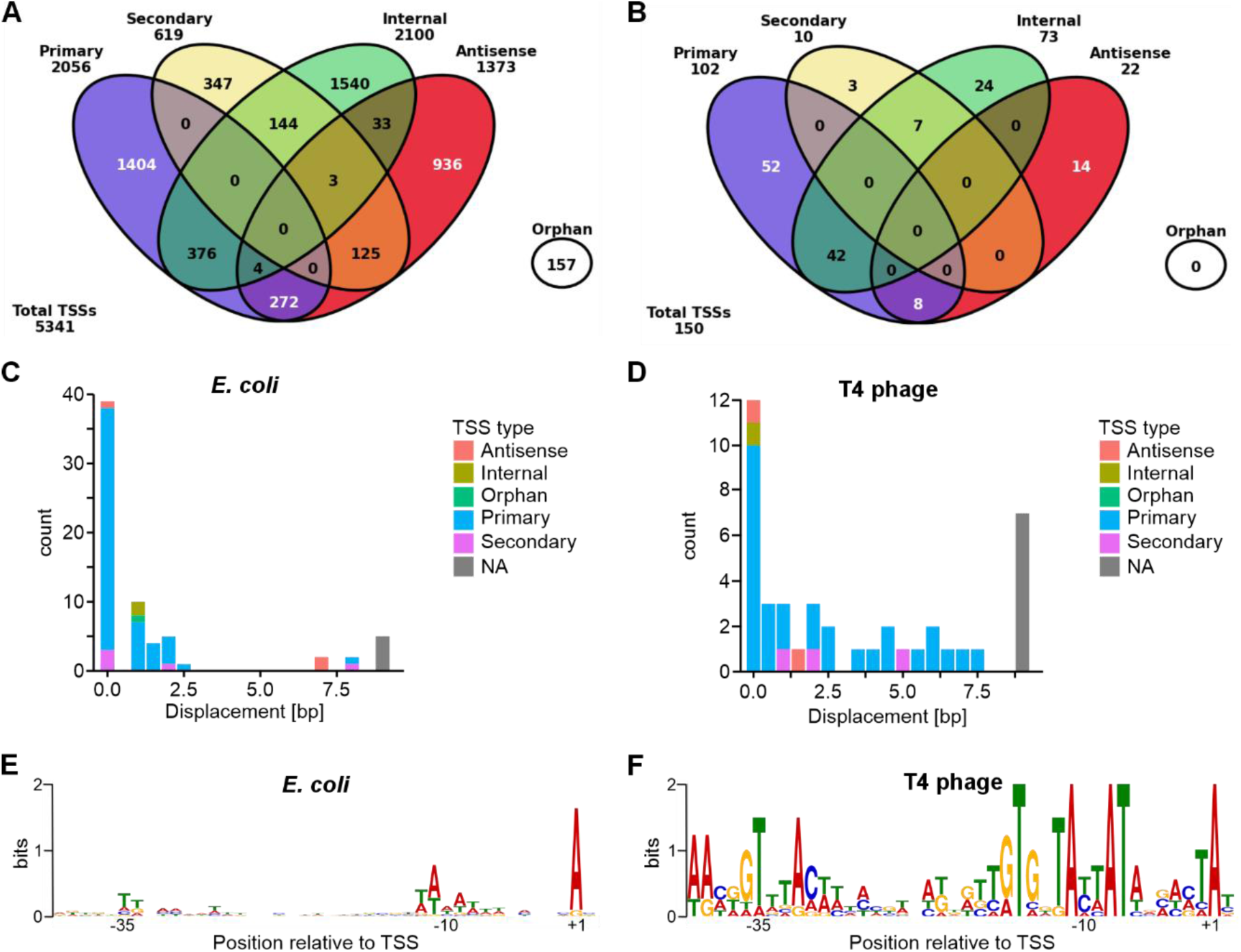
Global transcription start site (TSS) and NAD-RNA TSS analysis in *E. coli* and T4 phage. (**A**, **B**) Types and distribution of all TSS in *E. coli* (**A**) and T4 phage (**B**) during infection as revealed by time-resolved dual differential RNA-Seq (dRNA-Seq) during T4 phage infection of *E. coli* before (0 min) and 4, 10, 20 min post-infection (n=1). (**C**, **D**) Displacement of TSSs of NAD-RNAs identified by NAD captureSeq from “canonical” TSSs derived from dRNA-Seq data for *E. coli* (**C**) and T4 phage (**D**) NAD-RNAs from replicate 1. (**E**, **F**) Motifs of promoters of all NAD-RNAs of *E. coli* (42 promoters) (**E**) and T4 phage (32 promoters) (**F**) that were detected in both replicates (consensus). Motifs were created using Meme Suite (Bailey et al. 2015).

To reveal the regulatory logic behind the synthesis of NAD-capped transcripts, we set out to compare promoters and TSSs derived from dRNA-seq and NAD captureSeq. To the best of our knowledge, such a comparison is thus far unique. Therefore, we here use dRNA-seq-defined promoters or TSSs to distinguish those from NAD captureSeq-defined promoters and TSSs, which describe the 5’-ends of NAD-RNAs and their associated promoters. NAD captureSeq-defined NAD-RNA 5′-ends closely coincided with dRNA-seq-defined primary TSSs for both *E. coli* and T4 transcripts (Figure 4 C, D). The mean displacement of NAD captureSeq- to dRNA-seq-derived TSSs accounted for 1.6 bp (R1) or 1.4 bp (R2) for *E. coli* NAD-RNAs or 2.4 bp (R1) or 1.9 (R2) bp for T4 phage NAD-RNAs, respectively (Figure 4 C, D). Importantly, the TSSs of NAD-capped tRNA precursors (Supplementary Figure S7 A, B) agree with the TSSs of the corresponding tRNA primary transcripts without any displacement.

The *E. coli* NAD captureSeq-derived promoters share a +1A TSS (Figure 4E), which is an essential feature for *ab initio* NAD-capping of transcripts by the RNAP (Cahova et al. 2015; Bird et al. 2016), as well as an AT-rich −10 element and a T-rich −35 element. This pattern compares well to the overall *E. coli* +1A consensus promoter motif (Supplementary Figure S9A). It strikes out that *E. coli* NAD captureSeq-derived promoters have a preference towards C or T at the −1 position (Figure 4E). While this does not match the ideal bases (A or G) for NAD-capping at that position, it does agree with the finding that C and T most frequently occur at the −1 position in *E. coli* NAD-RNAs *in vivo* (Vvedenskaya et al. 2018).

The general consensus (dRNA-seq-derived) T4 phage promoter also contains an AT-rich −10 element and displays a tendency towards +1A TSSs (Supplementary Figure S9B) in agreement with the previously studied T4 promoter motif (Miller et al. 2003). 60 % (90 in total) of the T4 phage promoters identified by dRNA-seq initiate with adenosine. For 62 % of these promoters, we identified NAD-RNAs by NAD captureSeq. Similar as for *E. coli* NAD captureSeq-derived promoters, promoters associated to T4 NAD-RNAs share the +1A TSS and the AT-rich −10 element, too (Figure 4F). Interestingly, a comparison to the T4 consensus promoter indicates pronounced enrichment of thymine at −35 position and a TG motif downstream of the −10 element in the NAD captureSeq-derived promoters. The −335 element of strong promoters is typically AT-rich to enable the binding of the RNAP and efficient transcription (Sutherland and Murakami 2018). Importantly, the T4 NAD captureSeq-derived promoter features are not as enriched in the 90 T4 phage +1A promoters (Supplementary Figure S 9B). Thus, these distinct promoter elements may be involved in NAD-capping of the observed set of T4 phage NAD-RNAs.

Notably, the RNAP is successively modified during T4 phage infection to transcribe specific phage genes in the early, middle and late phases of infection. The RNAP is ADP-ribosylated early during infection (Koch et al. 1995; Wilkens et al. 1997), then associated with MotA and AsiA to drive middle transcription (Ouhammouch et al. 1995), followed by association with Gp55/Gp33 for late transcription (Nechaev et al. 2004). Gp55, for instance, is a diverged member of the sigma 70 family. Each of these forms of the RNAP recognizes distinct T4 promoters (Miller et al. 2003; Geiduschek and Kassavetis 2010; Hinton 2010). We thus analyzed the motifs of dRNA-seq and NAD captureSeq-derived promoters in each infection phase and recorded infection phase-specific sequence patterns (Supplementary Figure S9C). In early, middle and late promoters we found the characteristic −10 elements such as the early TGTG(A/G)TA(C/T)(A/T)AT motif (Miller et al. 2003; Hinton 2010), the middle TATAAT motif (Hinton 2010) or the TAAAT motif in late promoters (Geiduschek and Kassavetis 2010). Overall, this suggests that various forms of the *Ec*RNAP are capable of NAD-capping and that initiation of T4 transcription with NAD can occur at a diverse set of promoters during infection. Moreover, this emphasizes that both canonical, i.e. 5’-PPP-modified, and NAD-capped T4 phage transcripts are transcribed from the same TSSs and thus promoters by the *Ec*RNAP.

In conclusion, this hints at a possibly broader spectrum of regulatory potential of NAD-RNA synthesis than previously reported in *E. coli* (Cahova et al. 2015; Bird et al. 2016; Vvedenskaya et al. 2018) enabled by studying its infection by bacteriophage T4.

Finally, we used the comparison of dRNA-seq and NAD captureSeq data to more closely inspect the *dda.1* locus. The dRNA-seq- and NAD captureSeq-derived TSS perfectly overlap and this TSS represents the only detected TSS that could drive the expression of the entire locus spanning four genes (*dda.1* – *dexA.1*). The resulting untranslated region spans 101 nucleotides and contains an AG-rich motif 6 b upstream of the start codon, likely a Shine-Dalgarno sequence. To explain 35% of NAD-capping of the *dda.1* transcript (Figure 3B), we inspected the promoter more closely and identified that the −1 position is accommodated by adenine, which is known to drive higher degrees of NAD-capping (Vvedenskaya et al. 2018) allowing the RNAP to more efficiently initiate transcription with NAD. This could result in potentially more stable NAD-capped transcripts that would require degradation initiated by NAD-RNA decapping.

### The T4 phage Nudix hydrolase NudE.1 hydrolyses NAD-caps *in vitro*

We recorded both appearance and disappearance phage and host-derived NAD-RNAs during infection. This prompted us to investigate not only their synthesis but also potential pathways for their degradation. In *E. coli*, degradation of NAD-RNAs is initiated by NAD-RNA decapping by the Nudix hydrolase NudC (Cahova et al. 2015; Höfer et al. 2016b; Wolfram-Schauerte and Höfer 2023). We therefore asked whether additional Nudix hydrolases might contribute to NAD-RNA decapping during T4 phage infection. The T4 phage encodes a single Nudix hydrolase – termed NudE.1 – that has been described to hydrolyze adenosine-derived cofactors and metabolites, primarily ADP-ribose, Ap_3_A and FAD, within their pyrophosphate moiety (Xu et al. 2002). However, activity on NAD has not been reported (Xu et al. 2002).

We used Colabfold (Mirdita et al. 2022) to predict the structure of NudE.1, which reveals a wide-open cleft, in which the catalytically relevant Nudix motif is located (Figure 5A, Supplementary Figure S 10A, B). This architecture suggested that substrates such as NAD or NAD-RNA might also be accepted by NudE.1. In addition, the predicted NudE.1 structure aligned well to the *E. coli* Nudix hydrolase RppH (Supplementary Figure S10C), which trims the triphosphate 5’-terminus of primary transcripts (Deana et al. 2008). When modelled with a possible 10-nucleotide RNA substrate using Alphafold3 (Abramson et al. 2024), the terminal 5’-A nucleotide is positioned in close proximity to the catalytic Nudix motif suggesting a possible 5’-RNA processing activity for NudE.1 (Supplementary Figure S10D). On this basis, we speculated that NudE.1 may also accept substrates such as NAD and in particular NAD-RNA for hydrolysis. We expressed and purified NudE.1 migrating as an apparent monomer during size exclusion chromatography (Supplementary Figure S10E). Initially, we assessed the activity of NudE.1 on NAD using site-specifically ^32^P-labelled NAD. We observed that NudE.1 efficiently hydrolyses NAD into nicotinamide mononucleotide (NMN) and AMP, whilst a mutant of the catalytic site E64,65Q (NudE.1 E64,65Q) did not show hydrolysis activity on NAD (Figure 5B). Notably, NudC hydrolyses NAD less efficiently (0.56-fold rate constant of NudE.1) at the same substrate and enzyme concentrations *in vitro* (Figure 5B).

**Figure 5:**
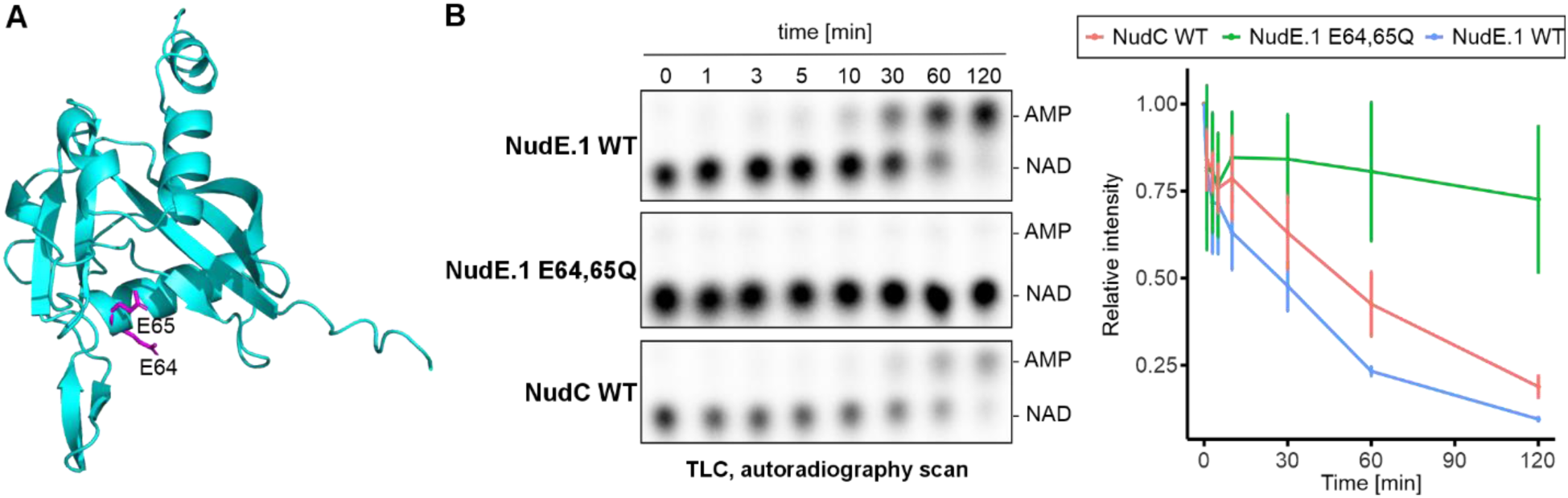
NAD hydrolysis by the T4 phage Nudix hydrolase NudE.1 *in vitro*. (**A**) Alphafold prediction for NudE.1 indicating E64 and E65 whose mutation to glutamines inactivates NudE.1 (E64,65Q). Prediction metrics are shown in Supplementary Figure S6A. (**B**) Activity of NudE.1 WT, NudE.1 E64,65Q and NudC WT on radiolabeled _32_P-NAD *in vitro* as monitored by thin-layer chromatography (TLC) and autoradiography. NudE.1 and NudC WT hydrolyze NAD giving rise to AMP and NMN. Error bars represent standard deviation from the mean, data points were each fitted with a generalized linear model (n=3).

Next, we set out to test the potential NAD-RNA decapping activity of NudE.1 on a 10-nucleotide NAD-RNA with a 3’-Cy5-label (NAD-10mer-Cy5) *in vitro* (Figure 6A). Similar to the known decapping enzyme NudC, NudE.1 efficiently (rate constant ∼9-fold higher than for NudC) decapped NAD-RNA *in vitro*, whilst the inactive mutant NudE.1 E64,65Q showed no measurable decapping activity (Figure 6B, Supplementary Figure S 11A). We then tested whether NudE.1 preferentially decaps distinct RNA structures at the 5’-NAD-cap but found no clear preference for different 5′-end structures (Figure 6C, Supplementary Figure S 11B). The Nudix hydrolase NudC has been reported to act preferentially on NAD-RNA rather than NAD due to its RNA binding affinity. Thus, *in vivo*, where NAD is far more abundant than NAD-RNA (approx. 700-fold molar excess) (Bennett et al. 2009; Chen et al. 2009; Wolfram-Schauerte et al. 2023), NudC hydrolyzes NAD-RNA rather than NAD (Höfer et al. 2016b). To mimic these conditions *in vitro*, we assessed NAD-RNA hydrolysis by NudE.1 and NudC in the absence and the presence of 700- and 1,500-fold molar excess of NAD over NAD-RNA. Notably, increasing NAD levels did not affect decapping by NudC (rate constant at 1,500-fold excess at 0.73 x of constant at 0-fold excess), whilst NAD-cap hydrolysis by NudE.1 was reduced (rate constant at 1,500-fold excess at 0.17 x of constant at 0-fold excess) (Figure 6D). However, under physiologically relevant ratios of NAD and NAD-RNA (700:1), NudE.1 still efficiently decapped NAD-RNA *in vitro*. Assessing the levels of NAD over the time course of the kinetic experiments, we barely saw any effects for both NudE.1 and NudC (Supplementary Figure S11C) indicating that both enzymes seem to exert an overall preference for NAD-RNA as substrate *in vitro*.

**Figure 6:**
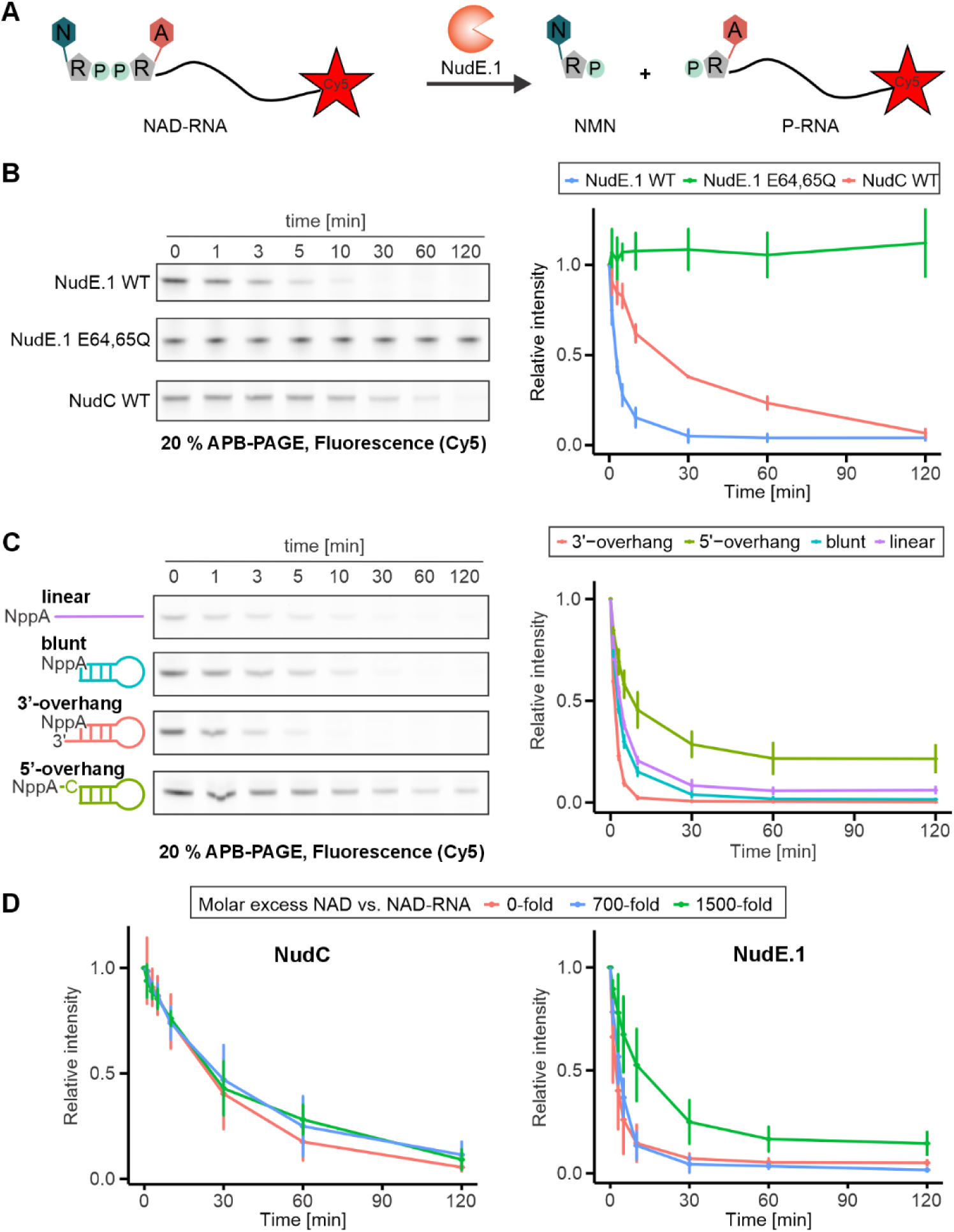
NAD-RNA decapping by the T4 phage Nudix hydrolase NudE.1 *in vitro*. (**A**) Reaction scheme for the NAD-RNA decapping by NudE.1 by hydrolysis of the pyrophosphate moiety in the NAD-cap. The resulting monophosphorylated RNA is separated from NAD-RNA during APB-PAGE. (**B**) NudE.1 WT efficiently decaps NAD-RNA *in vitro*. Cy5 scan visualizes NAD-RNA and 20 % APB-PAGE is employed to separate NAD- and PPP-RNA thereby indicating the specific decapping activity on NAD-RNA substrate (n=3). (**C**) Decapping activity dependence of NudE.1 on RNA substrate’s secondary structure analyzed by 20 % APB-PAGE and fluorescent Cy5 scan (n=3). (**D**) Effects of molar excess of NAD over NAD-RNA on the decapping activity of NudE.1 *in vitro* (n=3). Increasing the molar excess of NAD gradually decreases NudE.1 mediated NAD-RNA decapping (right panel), whilst decapping by NudC was barely affected (left panel). Error bars represent standard deviation from the mean, data points were each fitted with a generalized linear model (**B**, **C**, **D**; n=3). Corresponding large scale gel images for data in B and C are available (Supplementary Figure S 9A, B).

Given the evidence for potential RNA substrate binding by NudE.1 and its structural similarity to RppH (Supplementary Figure S 10B-D), we tested whether NudE.1 - similar to RppH - trims 5’-PPP-ends of RNA substrates. As expected, we only observed removal of the terminal ^32^P-labelled gamma-phosphate by RppH and neither for NudE.1 nor for NudC (Supplementary Figure S11D-F). Overall, these findings suggest NudE.1 as a Nudix hydrolase with NAD-RNA decapping activity *in vitro*. The predicted binding and positioning of the (NAD-)RNA substrate for decapping (Supplementary Figure S10A-D) could resemble a mechanistic model similar to the one reported for NudC (Höfer et al. 2016b), but rather in a monomeric state.

### Influence of NudE.1 on T4 phage infection of *E. coli*

To elucidate the functional role of NudE.1 *in vivo*, during T4 phage infection, we created a T4 phage with catalytically inactive NudE.1 (T4 NudE.1 E64,65Q) and characterized the phenotype of the latter (Pozhydaieva et al. 2024a). T4 NudE.1 E64,65Q displayed a lysis delayed significantly by approx. 15 minutes compared to T4 WT (Figure 7 A, B). Further, we detected similar amount of progeny released by T4 NudE.1 E64,65Q and T4 WT and similar burst sizes of both phages (Figure 7C, Supplementary Table S9). This is consistent with previous studies of a T4 phage lacking NudE.1, which suggested that NudE.1 plays an auxiliary rather than essential role during T4 phage infection (Xu et al. 2002; Miller et al. 2003).

**Figure 7:**
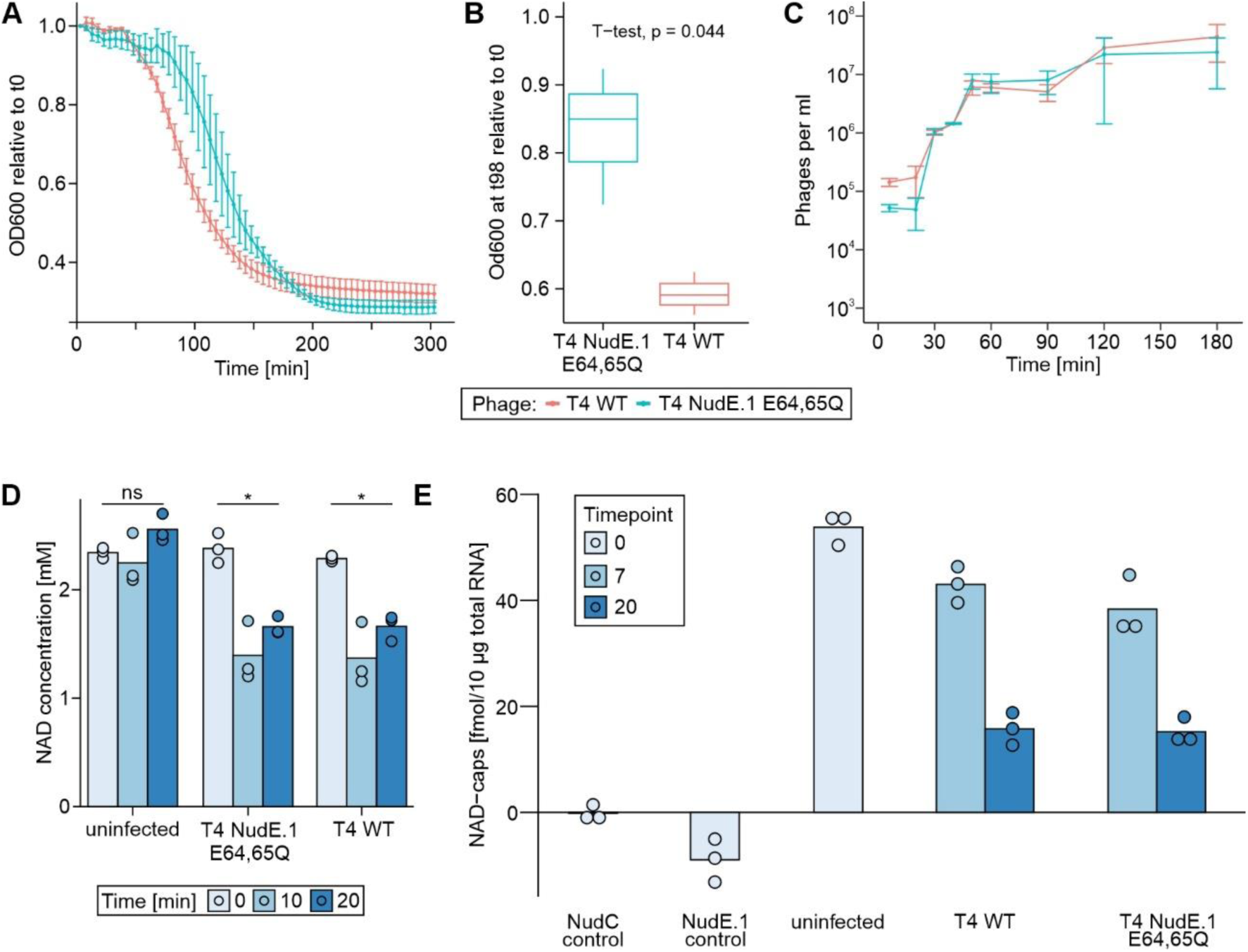
Phenotype of the NudE.1 E64,65Q mutant T4 phage compared to T4 phage WT. **(A)** Lysis curves of T4 phage WT and T4 phage NudE.1 E64,65Q (T4 NudE.1 E64,65Q) upon infection of *E. coli* B strain measured as OD_600_ over the time course of infection (n=3). OD_600_ values are normalized relative to the initial measurement at t0. **(B)** T-test comparing OD_600_ of T4 WT and T4 NudE.1 E64,65Q infected *E. coli* B strain at 98 minutes post infection given data presented in A (t-test, two-sided, p = 0.029 at p_signif_ < 0.05). **(C)** Progeny released from T4 WT and T4 NudE.1 E64,65Q infected *E. coli* measured as phages per mL over the time course of the burst size assay (n=3). **(D)** Same data as presented in Figure 1A complemented by measurements for T4 NudE.1 E64,65Q infection. Metabolomics analysis of NAD levels in *E. coli* infected with T4 WT, T4 NudE.1 E64,65Q or uninfected (none) over the time course of infection (n=3). NAD concentration [mM] refers to the concentration in an *E. coli* cell of 1 fL volume. **(E)** Same dataset as presented in Figure 1B, complemented by measurements obtained during infection with the T4 NudE.1 E64,65Q mutant and by a NudE.1 control. Total NAD-cap abundance was determined from RNA isolated from uninfected *E. coli* and from *E. coli* infected with either T4 WT or T4 NudE.1 E64,65Q for 10 min or 20 min. The NudC and NudE.1 controls represent total RNA from uninfected *E. coli* treated with recombinant NudC or NudE.1 prior to analysis and serve as negative controls, as enzymatic removal of NAD caps is expected to abolish the fluorescence signal. Consistent with this expectation, no detectable fluorescence signal was observed in either control. NAD-cap levels upon T4 NudE.1 E64,65Q infection were significantly reduced compared to uninfected *E. coli* (Welch’s two-sample *t*-test, *p* < 0.001), whereas no significant differences were detected between T4 WT and T4 NudE.1 E64,65Q infection. Bars represent mean values and dots indicate individual biological replicates (*n* = 3). Error bars represent standard deviation (s.d.).

To identify the influence of NudE.1 on cellular cofactors during T4 phage infection, we complemented our LC-MS-based metabolite measurements (Figure 1) with the T4 phage carrying inactive NudE.1. Comparing T4 WT and T4 NudE.1 E64,65Q infection, all three analyzed cofactors displayed similar trends (Figure 7D, Supplementary Figure S1). Consequently, one cannot conclude that NudE.1 specifically modulates cofactor pools, such as NAD, during T4 phage infection.

To investigate a potential role of NudE.1 in decapping NAD-RNAs during infection, we employed the FluorCapQ assay to determine NAD-cap levels in total RNA from *E. coli* infected with T4 NudE.1 E64,65Q, too. With our control treatment of *E. coli* total RNA with NudE.1 *in vitro*, we first confirmed its overall NAD-decapping activity (Figure 7E). Upon T4 NudE.1 E64,65Q infection, NAD-cap levels decrease over the time course of infection (Figure 7E) in a similar manner to that observed in T4 WT infected *E. coli* (Figure 1). Consequently, NudE.1 does not change the overall abundance of NAD-RNAs during infection. while we cannot exclude that NudE.1 might modulate the NAD-capping of specific transcripts, at least under the conditions tested in this study.

In conclusion, our findings indicate NudE.1 as the first phage-derived enzyme with robust NAD-RNA decapping activity *in vitro*. Although NudE.1 catalytic function is required for efficient phage infection under our experimental conditions, the E64,65Q substitution simultaneously abolishes activity towards multiple substrates, including NAD, NAD-RNA, FAD, ADP-ribose, and Ap_3_A. Consequently, the observed lysis delay cannot be attributed specifically to impaired NAD-RNA decapping, and future separation-of-function studies will be required to determine which substrate(s) underlie the *in vivo* requirement for NudE.1 catalysis.

## CONCLUSION AND OUTLOOK

Here, we present the first comprehensive description of the presence of NAD-capped RNAs during a phage–host interaction. While transcriptional reprogramming during infection has been extensively studied, the role of RNA 5′ end modifications remains largely unexplored. Using NAD captureSeq during T4 phage infection of *E. coli*, we identify NAD-RNAs derived from both the bacterial host and the infecting phage, thereby demonstrating that NAD-capping is not restricted to host transcripts but also occurs on phage RNAs. Importantly, we show that phage NAD-RNAs are synthesized from diverse promoters and are likely synthesized by distinct forms of the host RNAP across different stages of infection, indicating dynamic remodeling of the NAD-capped RNA landscape during T4 phage infection. In addition, we identify the T4 encoded Nudix hydrolase NudE.1 as a phage-derived enzyme capable of decapping NAD-RNA *in vitro*.

These findings may pinpoint an additional layer of RNA modification during phage infection and raise several key questions. In the following sections, we discuss: first, how phage NAD-RNAs are generated and which transcriptional features may promote NAD-capping; second, how phage-and host-encoded Nudix hydrolases may shape NAD-RNA stability through decapping; and third, how NAD-RNA metabolism might intersect with NAD-dependent defense and counter-defense mechanisms during phage-host interactions.

First, we considered the biosynthesis of NAD-capped RNAs during phage infection. The presence of co-existing phage and host NAD-RNAs extends the current view that phage infection primarily relies on transcriptional reprogramming at the level of promoter recognition and transcription factor recruitment (Tabib-Salazar et al. 2019). Rather than representing an additional layer of regulation per se, our findings suggest that RNA 5′-end modifications may contribute an additional and previously overlooked level of transcript diversity during infection. Our combined NAD captureSeq and TSS analyses indicate that both phage and host NAD-RNAs are synthesized by the *E. coli* RNAP, consistent with the fact that T4 relies entirely on host transcription machinery (Miller et al. 2003). In addition, NAD-RNA promoters display distinct sequence features that may favor NAD-dependent transcription initiation, most notably a strong preference for adenosine at the transcription start site.

Our promoter analyses further suggest that promoter architecture contributes to NAD-RNA formation. A substantial fraction of T4 promoters initiates with adenosine (90 of 150 promotors initiate with A), a known prerequisite for NAD incorporation as non-canonical initiating nucleotide (Cahova et al. 2015; Bird et al. 2016). These observations suggest that T4 promoter architecture favors NAD-capping by the host RNAP.

During T4 phage infection, however, transcription is extensively reprogrammed by phage-encoded proteins and post-translational modifications that alter the activity of the host RNAP (Miller et al. 2003; Tabib-Salazar et al. 2019). These include interactions with transcription factors such as AsiA, MotA, gp55 and gp3, as well as ADP-ribosylation by the phage enzymes Alt and ModA (Koch et al. 1995; Miller et al. 2003; Tiemann et al. 2004; Tabib-Salazar et al. 2019). Despite extensive phage-mediated remodeling of the host RNAP, NAD-RNAs are detected throughout early, middle, and late stages of infection, indicating that NAD-capped transcripts can be generated across distinct transcriptional phases of T4 infection.

A particularly striking example is the T4 transcript *dda.1*, which displays one of the highest NAD-capping levels observed in our dataset, with approximately 35 % of transcripts initiating with NAD rather than a canonical phosphorylated 5’end. Consistent with efficient NAD incorporation, the corresponding promoter initiates with adenosine at the transcription start site and perfectly matches the dRNA-seq-defined TSS. Notably, the promoter spans 101 bp, substantially longer than typical T4 promoters. This unusual architecture raises the possibility that additional, currently unknown factors may interact with the promoter region and influence transcription initiation or NAD-capping efficiency, an interesting question for future studies. Despite being an mRNA encoding the T4 helicase Dda, dda.1 remains highly abundant throughout infection, unlike other early transcripts such as motA (Wolfram-Schauerte et al. 2022). This raises the question of whether NAD-capping could contribute to this unusual stability and thereby ensure sustained production of a helicase essential for DNA replication during phage infection, which will require further investigation. Although the high NAD-capping efficiency of *dda.1* makes it an attractive candidate for functional analysis, the present study does not directly test whether NAD-capping contributes to its stability or expression during infection. The ability to generate NAD-capped transcripts may extend beyond the T4 system. While T4 relies entirely on host transcription machinery, many bacteriophages encode distinct single- and multi-subunit RNA polymerases (Basu and Murakami 2014; Sokolova et al. 2020). Notably, the RNAP of bacteriophage T7 has been shown to initiate transcription with cofactors such as NAD *in vitro*, suggesting that different transcription strategies may nevertheless converge on the production of cofactor-capped RNAs. NAD-capping may therefore represent a broader feature of phage transcriptomes rather than a phenomenon restricted to T4 infection (Huang 2003; Pozhydaieva et al. 2024b).

Moreover, given the diversity of non-canoncial RNA caps identified in bacteria, including FAD or UDP-GlcNAc-capped RNAs, (Julius and Yuzenkova 2017; Wang et al. 2019; Schauerte et al. 2021), additional cofactor-derived RNA modifications may likewise contribute to phage biology (Pozhydaieva et al. 2024b). Exploring their diversity, regulation and function therefore represents an important direction for future studies.

Secondly, we explored the possible role of NAD-RNA decapping during phage infection. NAD-RNA decay is typically initiated by decapping through Nudix hydrolases such as NudC, representing an important mechanism for regulating RNA stability in bacteria (Höfer et al. 2016; Cahová et al. 2015). In this context, we identify NudE.1 as the first phage-encoded enzyme with NAD-RNA decapping activity *in vitro*. This finding suggests that T4 may not only influence the synthesis of NAD-capped RNAs through transcriptional reprogramming but may also directly modulate their turnover by the presence of its own decapping enzymes.

Although NudE.1 does not exert a dominant effect on the global NAD or NAD-RNA pool under the tested conditions, proteomics data show that it is up to 60-fold more abundant than the host enzyme NudC during infection (Wolfram-Schauerte et al. 2022). This suggests that NAD-RNA stability may be shaped by the combined activities of host- and phage-encoded decapping enzymes and raises the possibility that T4 partially redirects NAD-RNA metabolism toward phage-controlled pathways (Xu et al. 2002).

More broadly, selective NAD-RNA decapping could provide a mechanism to influence transcript stability and gene expression during infection. However, because NudE.1 acts on multiple nucleotide-derived substrates, the specific contribution of NAD-RNA decapping to phage replication, host adaptation, and defense or counter-defense processes remains to be determined (Wang et al. 2023; Osterman et al. 2024a). The physiological RNA substrates of NudE.1 remain unknown. Future transcript-resolved analyses will be required to determine whether NudE.1 acts on specific host or phage NAD-capped RNAs during infection.

Finally, Nudix hydrolases are not unique to T4 but are also encoded by diverse phages (Lee et al. 2017). For example, the *Vibrio* phage KVP40 employs a Nudix hydrolase in an NAD salvage pathway, highlighting the broader relevance of this enzyme class in phage biology (Lee et al. 2017). Consistent with this, homologues of NudE.1 can be found in various phages targeting *Escherichia*, *Salmonella* and *Shigella* (Supplementary Table S10). Together, these findings suggest that Nudix hydrolases may represent a conserved strategy to modulate cofactor metabolism and RNA stability during phage infection.

Finally, we considered how NAD and NAD-capped RNAs may contribute to regulatory processes during phage infection. Our findings suggest that NAD-RNAs may function at multiple interconnected levels during infection, including as substrates for post-translational modifications such as RNAylation, as modulators of cellular NAD homeostasis through decapping and recycling pathways, and, consequently as factors influencing NAD-dependent defense and counter-defense mechanisms during infection. In particular, RNAylation represents an important functional link between NAD-capped RNAs and phage biology. Recent work demonstrated that the T4 ADP-ribosyltransferase ModB can transfer NAD-capped RNAs onto host proteins, thereby establishing RNAylation as a novel post-translational modification pathway during infection (Wolfram-Schauerte et al. 2023). Notably, several host and phage transcripts identified as RNAylation substrates overlap with NAD-RNAs detected in our study, supporting a functional role of NAD-capped RNAs beyond transcription initiation (Wolfram-Schauerte et al. 2023). Notably, only a subset of NAD-capped RNAs identified to date has been reported to serve as RNAylation substrates. Because NAD-capped RNAs are substantially more widespread than known RNAylation substrates, the determinants of RNAylation specificity remain unclear and warrant further investigation. A second potential regulatory layer may arise from the coupling between NAD-RNA metabolism and cellular NAD homeostasis. NAD-capped RNAs persist despite a progressive decline in intracellular NAD levels (Wiedermannova et al. 2024). Importantly, while changes in NAD metabolism have previously been investigated during bacteriophage T7 infection, our data provide a direct comparison of intracellular NAD and NAD-RNA levels throughout T4 infection (Ofir et al. 2021; Wiedermannova et al. 2024). We observe parallel declines of intracellular NAD and NAD-RNA levels during infection, consistent with a relationship between cofactor availability and NAD-RNA formation.

In recent years, several bacterial defense systems have been shown to protect against phage infection by depleting or transforming NAD, highlighting NAD metabolism as a central battleground in phage–host interactions (Ofir et al. 2021; Garb et al. 2022; Zaremba et al. 2022; Osterman et al. 2024b). In this context, NAD is not only a metabolic cofactor but also a molecule that can be actively manipulated by both host and phage to regulate the course of infection. Consistent with this concept, NAD-capped RNAs in eukaryotic systems have been shown to be processed by the 5′–3′ exoribonuclease XRN1, directly linking RNA decay to NAD turnover and recycling (Sharma et al. 2022). By analogy, similar mechanisms may operate during phage infection, where the decapping of NAD-RNAs could couple RNA metabolism to cofactor homeostasis. During T4 phage infection, for example, NAD-RNA decapping could contribute to NAD recycling through the release of intermediates such as NMN. Taken together, our findings suggest that NAD and NAD-RNA are integrated into a broader regulatory network connecting RNA processing, metabolism, cofactor homeostasis and host-phage interactions. Mechanisms that alter intracellular NAD availability may therefore simultaneously affect NAD-RNA formation and function, either indirectly through cofactor availability or directly through NAD-RNA processing pathways, thereby contributing to infection-associated regulatory processes.

Importantly, the present study establishes the molecular inventory and temporal dynamics of NAD-capped RNAs during T4 infection, providing a descriptive framework for future functional studies. While our data demonstrate extensive remodeling of the NAD-RNA pool during infection, they do not directly address whether NAD-capping causally influences transcript stability, translation, or infection outcome. Future experiments employing targeted perturbation of NAD-cap formation, for example through transcription start site substitutions that reduce NAD incorporation, will be required to distinguish regulated NAD-capping from a passive consequence of promoter architecture and cellular NAD availability.

Overall, this study provides the first identification of NAD-capped RNAs in a bacteriophage and demonstrates their dynamic regulation during infection. These findings establish RNA 5′-end modifications as a previously unexplored layer of phage–host interaction biology and suggest that RNA-based regulatory strategies may play important roles in prokaryotic virus–host interactions. Given that cofactor-capping in eukaryotic viruses can contribute to immune evasion (Sherwood et al. 2023), analogous RNA-based regulatory strategies may likewise play important and previously unrecognized roles in prokaryotic virus–host interactions.

## Data Availability

Custom scripts, analyses and associated data are available under https://github.com/MaikTungsten/PhageEpitranscriptomics.

Differential RNA-Seq data is deposited in GEO database under accession GSE255091 (reviewer token: kdupuycmjdcvzwx). Nanopore sequencing data from NAD captureSeq is available from SRA under Bio Project PRJNA1073512 (reviewer access: https://dataview.ncbi.nlm.nih.gov/object/PRJNA1073512?reviewer=vup80knotd2d8tertrscl4nlp3). Nanopore data for T4 phage NudE.1 E64,65Q mutant screening is deposited in SRA under Bio Project PRJNA1075486 (reviewer access: https://dataview.ncbi.nlm.nih.gov/object/PRJNA1075486?reviewer=qv3g5fm72dn5r3t6qajph3lb34).

Custom scripts, analyses and associated data are available under https://github.com/MaikTungsten/PhageEpitranscriptomics. Differential RNA-Seq data is deposited in GEO database under accession GSE255091. Nanopore sequencing data from NAD captureSeq is available from SRA under Bio Project PRJNA1073512. Nanopore data for T4 phage NudE.1 E64,65Q mutant screening is deposited in SRA under Bio Project PRJNA1075486.

## SUPPLEMENTARY DATA

Supplementary Data are available at NAR online.

## AUTHOR CONTRIBUTIONS

M.W.-S., H.K. and A.M. performed total RNA isolations, established and performed NAD captureSeq, performed qPCRs to validate sequencing data. M.W.-S. cloned, expressed, purified and characterized NudE.1. M.W.-S. prepared samples for metabolomics, proteomics. M.W.-S. and H.K. performed experiments to validate NAD-RNAs. H.K. established the CapPro Assay. N.Po. performed T4 phage mutagenesis and screening PCR, A.A.R.R. performed barcoding and Nanopore sequencing, supervised by D.S. M.W.-S. analysed all sequencing data. M.W.-S. characterized the T4 phage mutant. M.W.-S. prepared samples for LC-MS/MS, N.Pa. performed metabolomics measurements, N.Pa. and M.W-S. analysed the data. S.K. performed quantitative nucleoside MS analysis. H.K. established and applied the FluorCapQ Assay. K.H. supervised the work. M.W.-S. and K.H. designed the study and wrote the first draft of the manuscript. All authors contributed to editing and proof-reading.

## ACKNOWLEDGEMENTS

We thank Peter Claus, Petra Mann and Ahmet Sanal for experimental assistance and the entire team of the Höfer lab and collaborators for fruitful discussions and the nice working environment. We thank the anonymous reviewers for their helpful comments towards improving this manuscript.

