## Supplementary files for "T4 phage RNA is NAD-capped and alters the NAD-cap epitranscriptome of *Escherichia coli* during infection"

**Content:**

Supporting Figures S1-S10

Supporting Tables S1-S5

Captions for separate Supporting Tables S6-S10

Supporting references

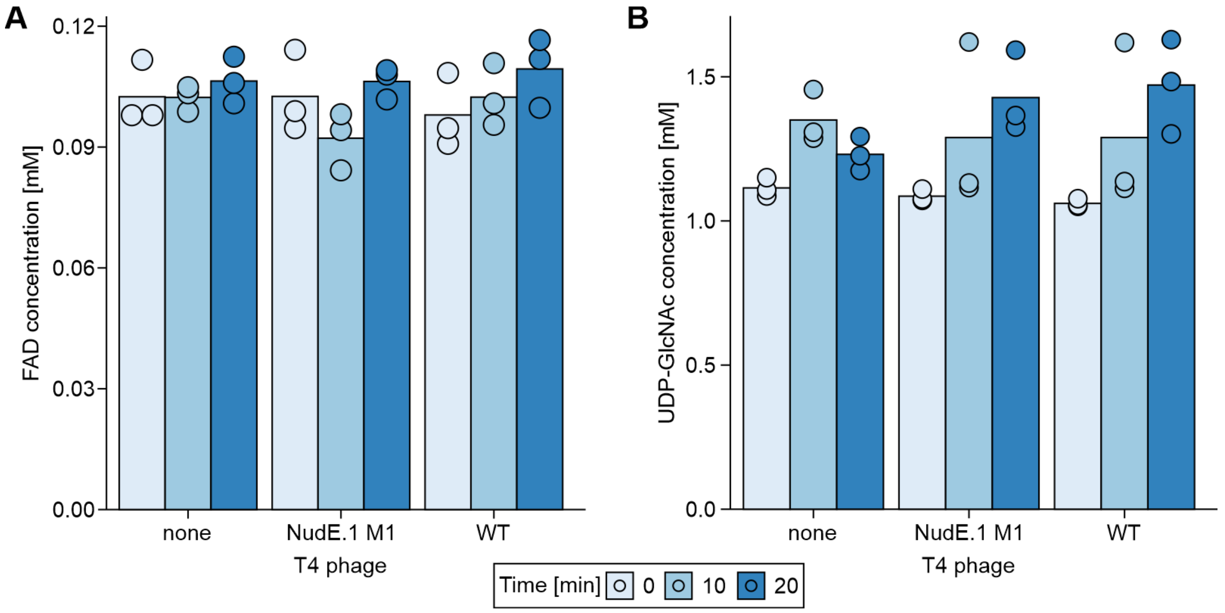

**Supplementary Figure S1: FAD and UDP-GlcNAc concentrations in during T4 phage infection.** Cellular concentrations of FAD (**A**) and UDP-GlcNAc (**B**) in T4 WT, T4 NudE.1 E64,65Q infected and uninfected (none) *E. coli* over the time course of 20 minutes before infection (0 min) as well as 10 and 20 min post infection. Concentrations are derived from endometabolomics analysis of biological triplicates (n=3) by LC-MS.

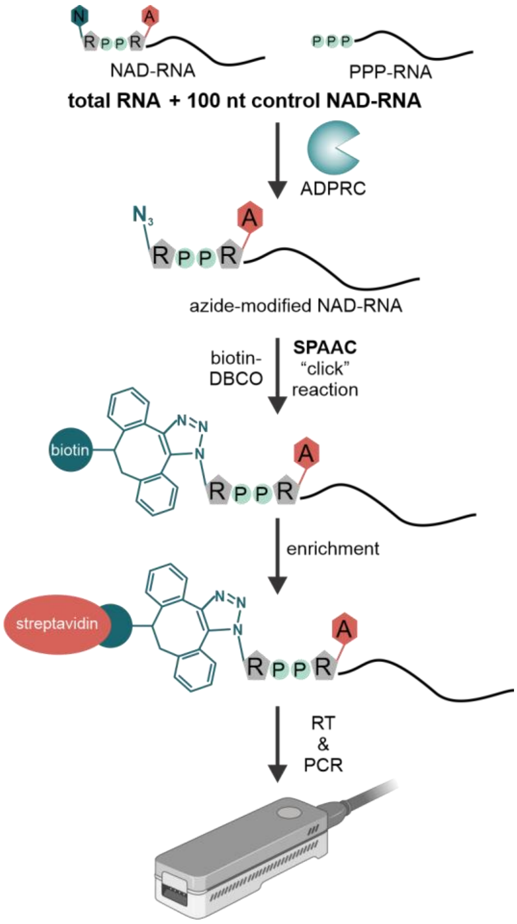

**Supplementary Figure S2: Adjusted NAD captureSeq workflow for the detection of NAD-capped RNAs.**

Schematic illustration of the optimized NAD captureSeq workflow. Total RNA with a spike-in of a 100 nt control NAD-RNA is subjected to treatment with ADP-ribosyl cyclase (ADPRC) using 3-azido-propanol as substrate yielding azide-modified NAD-RNAs. These are subsequently linked to biotin-dibenzocyclooctyne (DBCO-biotin) using strain-promoted azide-alkyne cycloaddition (SPAAC) (Hu et al. 2021; Zhang et al. 2021). Biotin-modified RNAs are captured on a streptavidin resin, reverse transcribed, PCR amplified and finally sequenced in a multiplexed fashion on a Nanopore Flongle flow cell. MinION schematic created with Biorender.com.

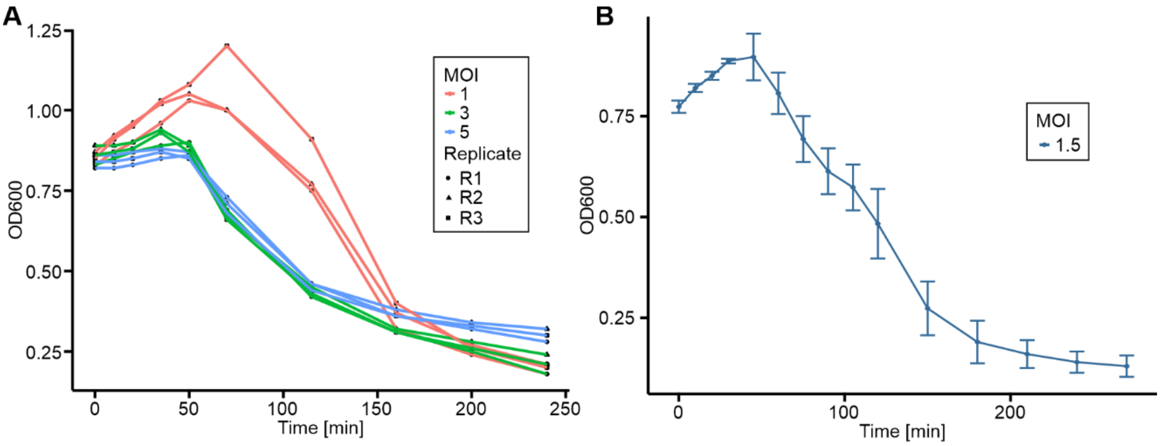

**Supplementary Figure S3: Lysis curves of T4 phage infection depending on MOI.**
(**A**) Lysis curves of *E. coli* upon T4 phage infection at multiplicity of infection (MOI) of 1, 3 and 5 measured by optical density at 600 nm ($OD_{600}$) (n=3). (**B**) Lysis curve for *E. coli* infected by T4 phage at an MOI of 1.5, the condition used for NAD captureSeq experiments. Data points with error bars represent mean ± s.d. (n=3).

| | **Replicate 1** | **Replicate 2** |
|---|---|---|
| Pass reads | 220,028 | 499,383 |
| Median read length [b] | 324 | 318 |
| Median PHRED score | 10.239 | 10.796 |
| Mean reads per barcode | 14,777 | 36,255 |
| Identified NAD-RNAs | 116 | 110 |

**Supplementary Figure S4: Statistics of Nanopore sequencing runs for both biological replicates presented in this study.**

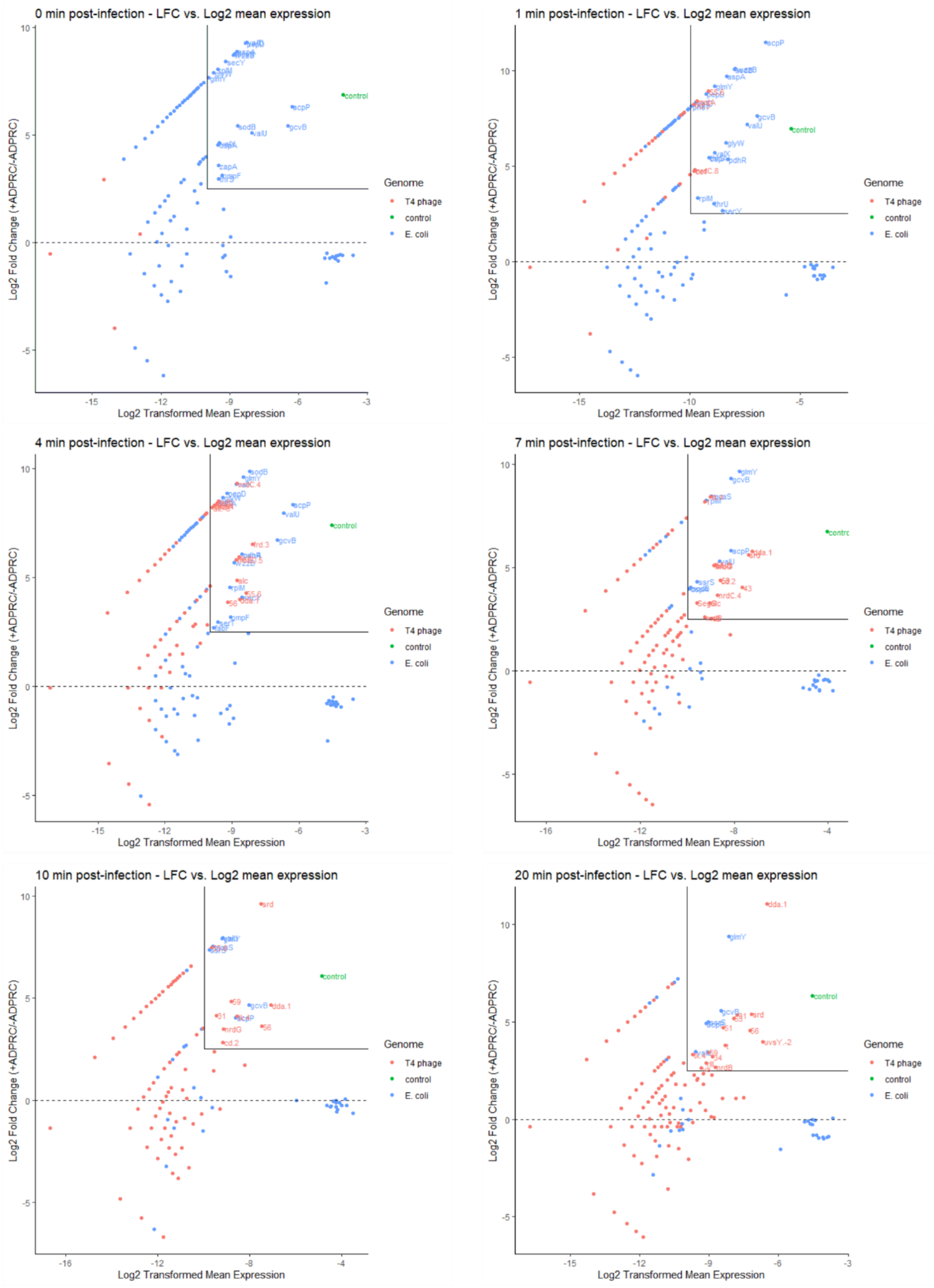

**Supplementary Figure S5: NAD captureSeq analysis for all time points for replicate 1.**
MA plots are presented for all 6 time points of infection (t0, t1, t4, t7, t10, t20) for replicate 1. y-axis represents the log2 fold change in normalized read counts comparing fully-treated sample (+ADPRC) and negative control (-ADPRC), x-axis shows log2 transformed mean normalized read counts for genes from + and -ADPRC samples. Enriched genes are labelled with their corresponding gene symbol and colored according to their genome (T4 phage, red; 100 nt control RNA, green; *E. coli*, blue).

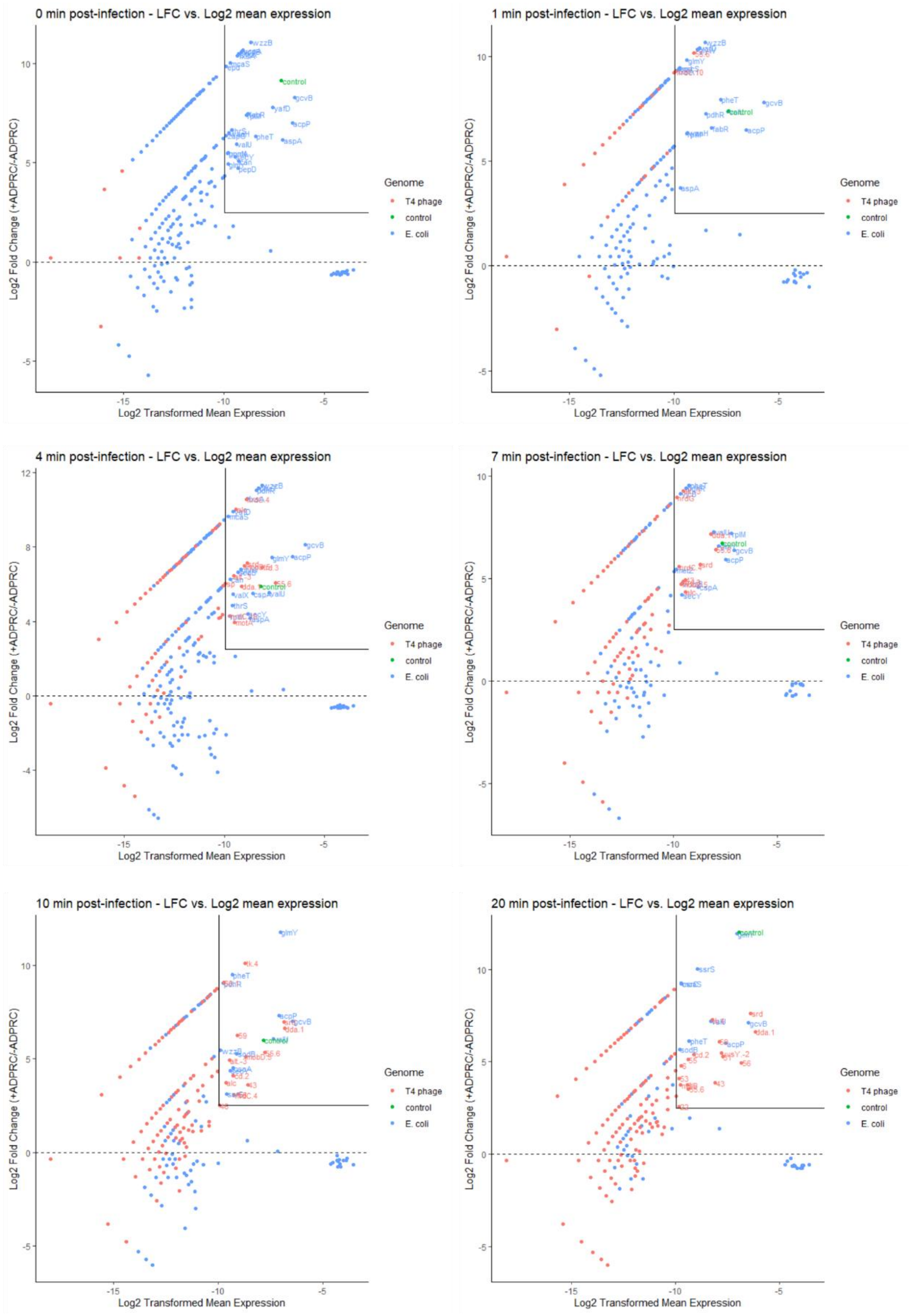

**Supplementary Figure S6: NAD captureSeq analysis for all time points for replicate 2.**
MA plots are presented for all six time points of infection (t0, t1, t4, t7, t10, t20) for replicate 2. y-axis represents the log2 fold change in normalized read counts comparing fully-treated sample (+ADPRC) and negative control (-ADPRC), x-axis shows log2 transformed mean normalized read counts for genes from + and -ADPRC samples. Enriched genes are labelled with their corresponding gene symbol and coloured according to their genome (T4 phage, red; 100 nt control RNA, green; *E. coli*, blue).

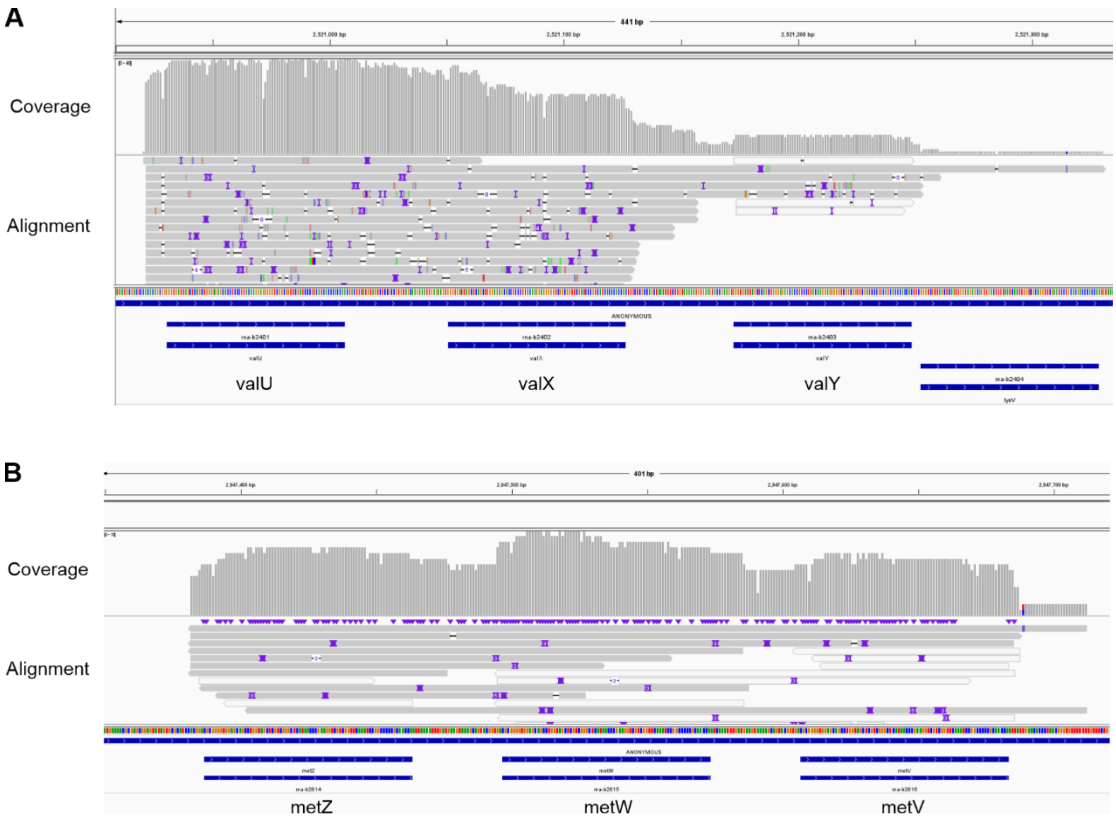

**Supplementary Figure S7: tRNA coverage in NAD captureSeq data.**
Coverage and alignment profiles for the valU/X/Y (**A**) and metZ/W/V (**B**) operons for the +ADPRC sample from time point t0, replicate 1 from the NAD captureSeq experiment. Reads clearly span across two or three tRNA genes in these operons indicating that the polycistronic tRNA precursors are NAD-capped in *E. coli*. The TSS is in good agreement with the TSS of primary transcripts derived from our dRNA-Seq data.

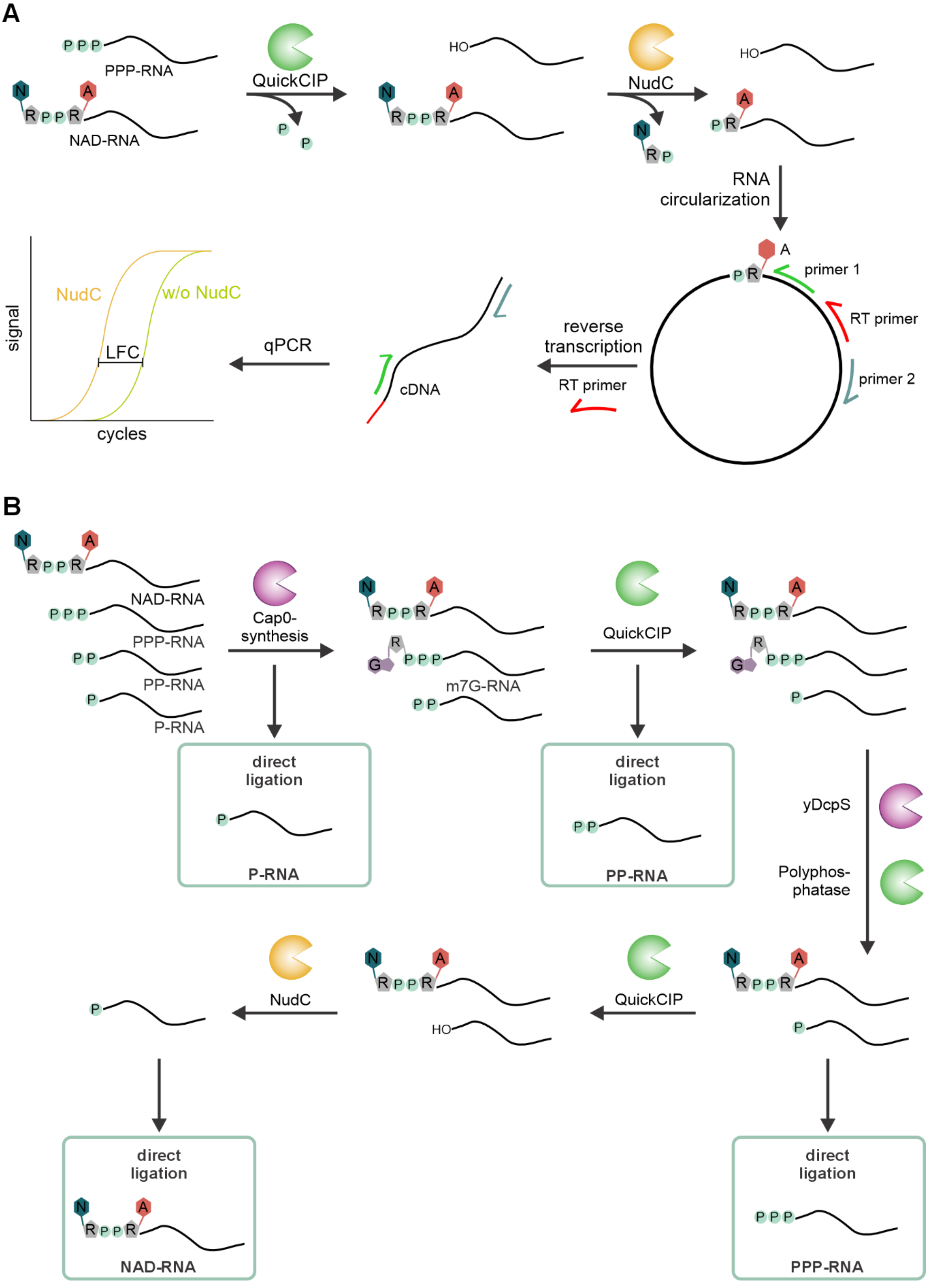

**Supplementary Figure S8: Validation of NAD-RNAs via circNC-like assay and quantitative profiling of 5´- end modifications (PPP, P, NAD) via CapPro assay**

(**A**) Ligation-based assay to prove the existence of specific NAD-RNAs. Briefly, total RNA is subjected to QuickCIP-treatment followed by NudC digest leaving a monophosphate only on formerly NAD-capped RNAs. RNA is then circularized by ligation and reverse transcribed with a target specific RT primer creating a unique cDNA spanning over the +1A position. The cDNA serves as a template for qPCR with target-specific primers to assess cDNA abundance in a NudC-treated and control sample as log2 fold change (LFC). (**B**) Schematic overview of the CapPro assay workflow. Total RNA is treated with sequential enzymatic steps to distinguish and quantify distinct 5′ end modifications. Therefore, RNA is treated with specific combinations of decapping or modifying enzymes: (i) NudC to selectively detect NAD caps, (ii) vaccinia capping enzyme and yDcpS to selectively detect PPP-RNA, or (iii) alkaline Phosphatase to selectively detect PP-RNAs. Subsequently, all RNAs are ligated using a circularization-based approach that selectively amplifies RNAs with accessible 5′P ends. qPCR is then used to quantify specific transcripts across treatments, allowing determination of the relative proportions of 5′-NAD, 5′-PPP, and 5′-P ends.

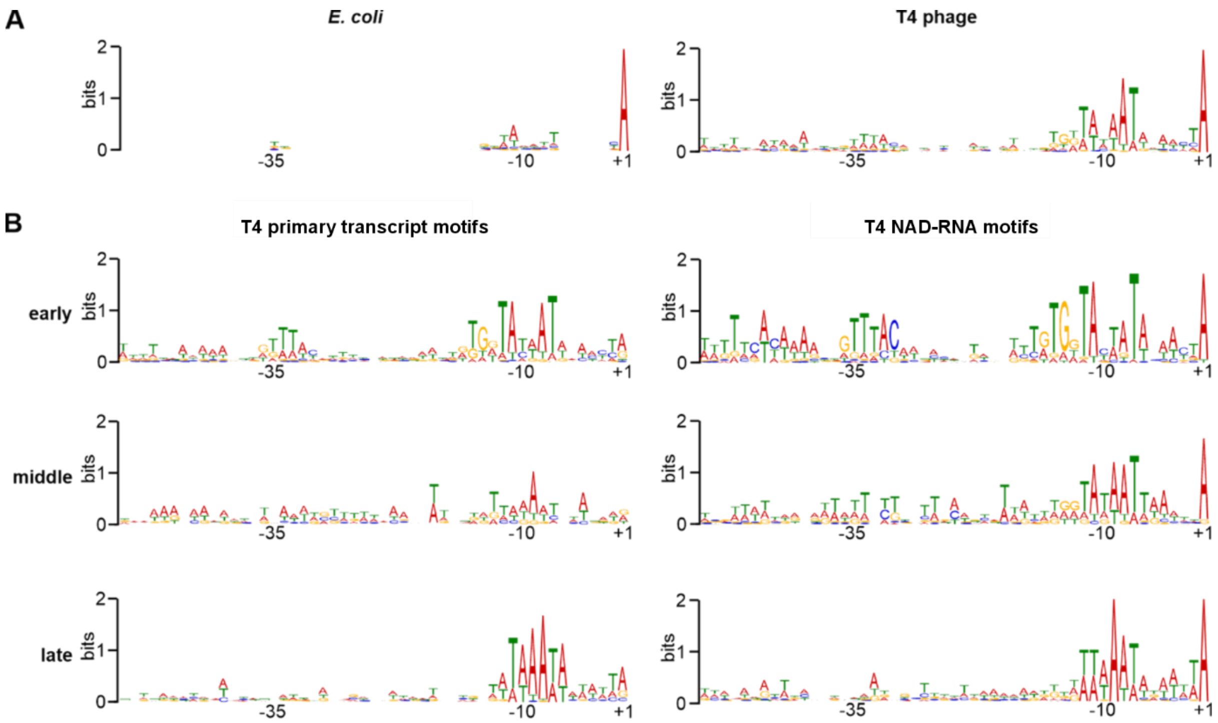

**Supplementary Figure S9: Sequence motifs associated with +1A transcription start sites and infection phase-specific T4 phage transcripts.**

(**A, B**) Sequence motifs derived from promoter regions upstream of +1A transcription start sites (TSSs) identified by differential RNA-seq (dRNA-seq) in *E. coli* (2081 TSSs, left) and T4 phage (90 TSSs, right). (**B**) Sequence motifs associated with infection phase-specific T4 primary transcripts identified by dRNA-seq (left) and infection phase-specific T4 NAD-RNAs identified by NAD captureSeq (right). Motifs were generated from 73 (dRNA-seq) or 24 (NAD-RNA) early, 23 or 17 middle, and 47 or 20 late transcript-associated TSSs, respectively. The transcription start site (+1) as well as the −10 and −35 promoter elements are indicated for each motif. All motifs were generated using MEME Suite (Bailey et al. 2015) from promoter regions spanning 51 bp upstream and downstream of the corresponding TSSs.

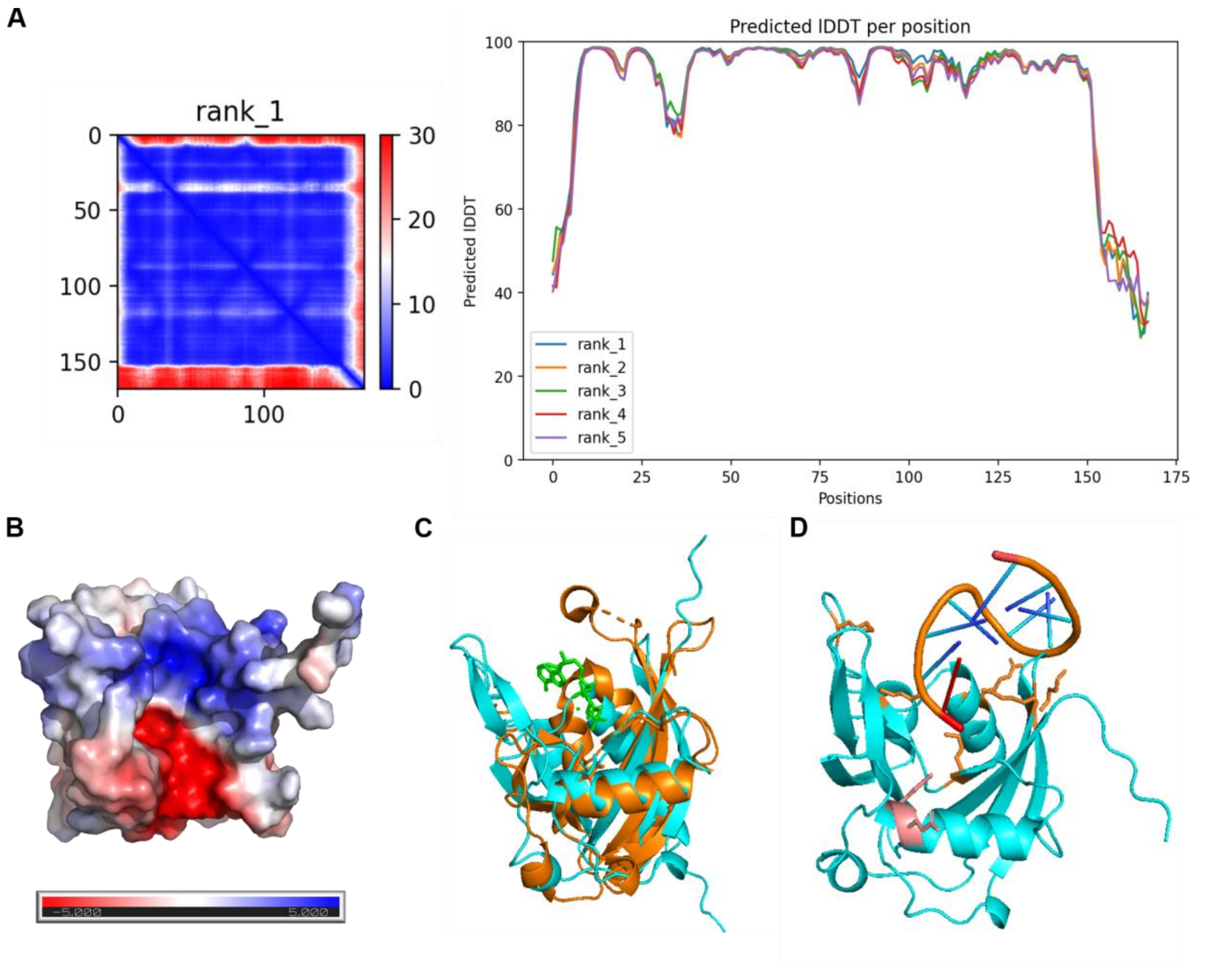

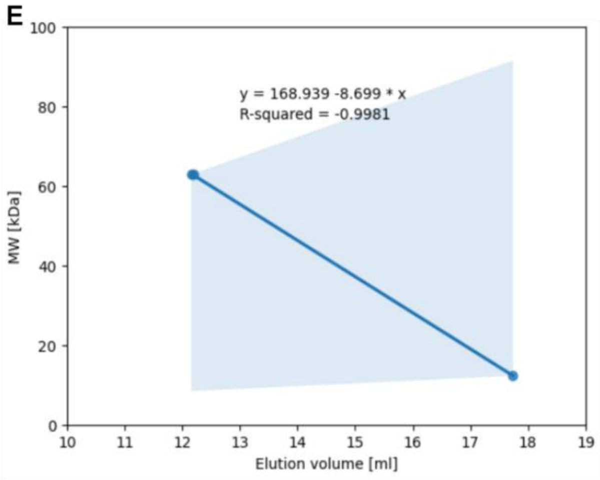

| | NudE.1 WT | NudE.1 M1 |
|---|---|---|
| Molecular weight [kDa] | 19.68 | 19.86 |
| Observed elution volume [ml] | 17.2 | 17.3 |
| Expected elution volume [ml] | 17.1 | 17.2 |
| Probable oligomeric state | monomer | monomer |

**Supplementary Figure S10: Structure and oligomeric state of the Nudix hydrolase NudE.1.**
(**A**) Alphafold prediction metrics for the Alphafold model of NudE.1 WT presented in Figure 6A. The model presented corresponds to rank 1. (**B**) Surface charge structural model of NudE.1 WT based on the model presented in Figure 6A. Red color indicates negative charge, blue color represents positive charge. Structural model enables a view directly in the open cleft with the catalytic site (charged negatively, colored red). (**C**) Structural alignment of NudE.1 model and a crystal structure of RppH with ATP (7PS3) with an RMSD of 1.636 Å. Ions have been removed from the structure, ATP shown in green, NudE.1 in cyan, RppH in orange. The two neighboring glutamate residues of the Nudix motifs of RppH and NudE.1 are shown and well align in the given alignment pointing towards the ATP/substrate binding site. (**D**) Structural model of NudE.1 along with the 10mer RNA used in this study generated by Alphafold3. E64 and E65 are shown as side chains in salmon, the 5′-adenosine of the RNA is highlighted in red and is positioned in the catalytic site. Selected arginine and lysine residues of NudE.1, which could interact with the RNA backbone, are shown as side chains in orange. (**E**) Analytical size exclusion chromatography

(SEC) to determine the oligomeric state of NudE.1. The SEC column was calibrated with monomeric protein standards of known molecular weight and a linear regression model was fit to calculate an expected elution volume for a given molecular weight. NudE.1 WT and E64,65Q migrate as apparent monomers during SEC.

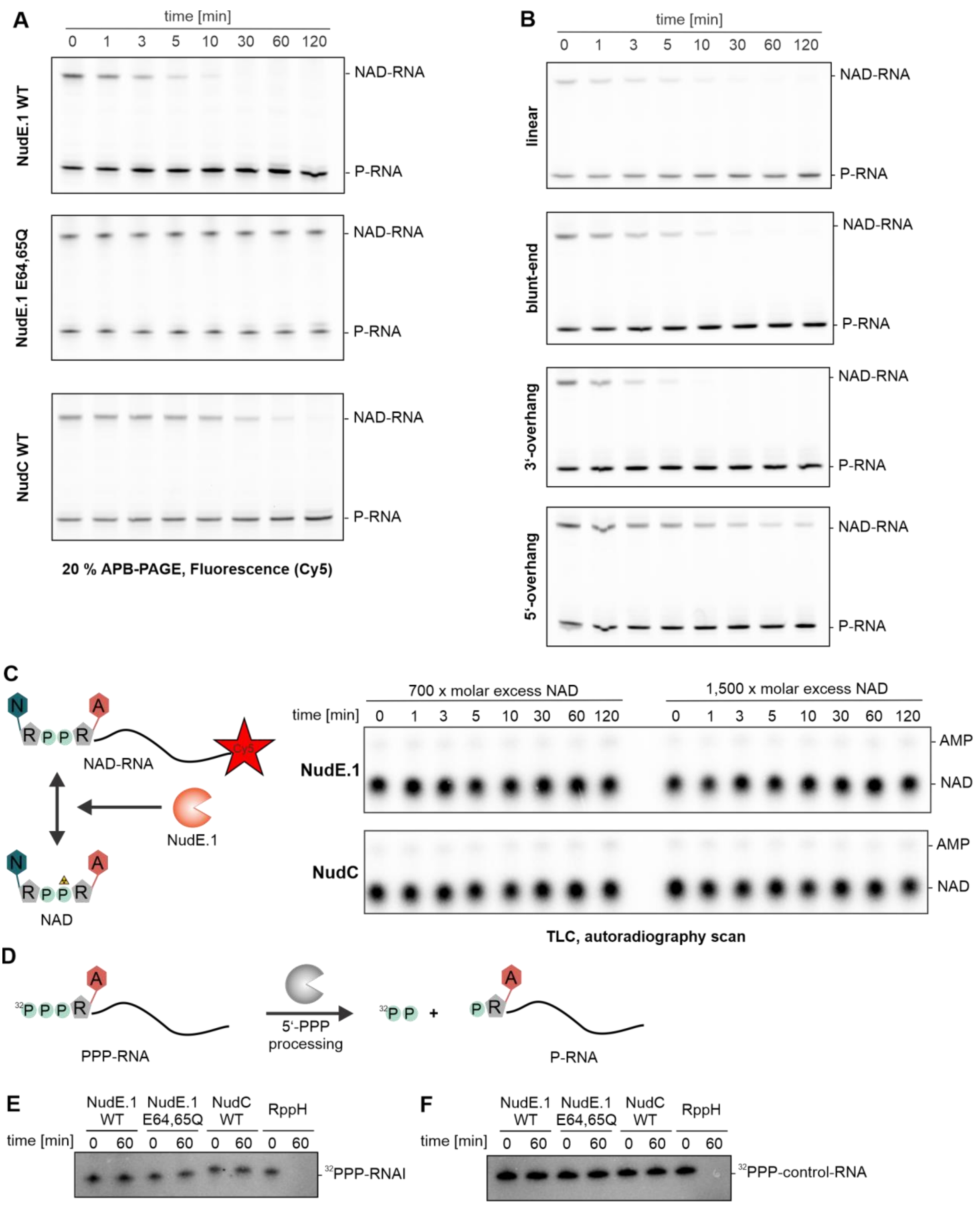

**Supplementary Figure S11: Processing activities by Nudix hydrolase NudE.1 *in vitro*.**
(**A**, **B**) Gel sections corresponding to gel segments of NAD-RNA bands presented in Fig. 6B,C, which additionally show the 5'-P-RNA band whose intensity increases upon NAD-RNA decapping by Nudix hydrolases NudE.1 or NudC, respectively. (**C**) NAD levels in the NAD vs. NAD-RNA competition assays as shown in Figure 6E. Autoradiography image of TLC analysis is shown. NAD levels are barely affected over

the time course of infection in the presence of both NudE.1 WT and NudC WT independent of the fold molar excess of NAD over NAD-RNA. (**D**) Schematic representation of enzymatic 5′-PPP to 5′-P/PP trimming, where the RNA is radiolabelled through $^{32}P$ at the gamma-phosphate, which is removed from the RNA upon trimming to 5′-P/PP-RNA. (**E**, **F**) 10 % PAGE analysis of processing of 5′-PPP-RNA to 5′-P/PP-RNA by Nudix hydrolases. Reduction of radioactive signal indicates processing to 5′-P-RNA. Analysis is shown for $^{32}$PPP-RNAI (E) and $^{32}$PPP-control-RNA (F) as substrates using NudE.1 WT, NudE.1 E64,65Q, NudC WT and RppH (as positive control) (n=1 each). Only RppH was found to remove the terminal gamma-phosphate.

**Supplementary Table S1: List of bacteria and phages used in this study.**

| Strain | Purpose | Reference |
|---|---|---|
| *E. coli* BL21 DE3 NudE.1 WT | Overexpression of His-tagged NudE.1 WT via pET28-NudE.1 WT | This study |
| *E. coli* BL21 DE3 NudE.1 E64,65Q | Overexpression of His-tagged NudE.1 M1 via pET28-NudE.1 E64,65Q | This study |
| *E. coli* BL21 DE3 NudC WT | Overexpression of His-tagged NudC WT via pET28-NudC WT | (Höfer et al. 2016) |
| *E. coli* B strain | Strain used for T4 phage infection | DSMZ (DSM 613) |
| *E. coli* JM109 + pUC19 | *E. coli* strain with NAD-capped RNAI | (Cahova et al. 2015) |
| *E. coli* BL21 DE3 Cas13a_NudE.1 | *E. coli* strain expressing Cas13a targeting NudE.1 WT RNA. | This study |
| *E. coli BL21 DE3* pET28a_NgTET_NudE.1-M1 + pCpf1_NudE.1 | *E. coli* expressing NgTET and Cas12 targeting NudE.1 WT, and providing NudE.1 M1 donor DNA | This study |
| Bacteriophage T4 WT | WT T4 phage | DSMZ (DSM 16352) |
| Bacteriophage T4 NudE.1 E64,65Q | T4 phage with NudE.1 E64,65Q (inactive) | This study |

**Supplementary Table S2: DNA sequences of probes, primers and splints.**

| Name | Sequence |
|---|---|
| **Northern blot probes** | |
| Northern probe RNAI | TGTTTGCCGGATCAAGAGCTACCAACTCTTTTTCCGAAGGTAACTGGCTTCAGCAGAGCGCAGATACCAAATACTGT |
| Northern probe 5S rRNA | CCCCACACTACCATCGGCGCTACGGCGTTTCACTTCTGAGTTCGGCATGGGGTCAGGTGGGACCACCGCGCTACGGC |
| **qPCR primers for NAD captureSeq data validation** | |
| qPCR fwd primer 100 nt control RNA | ATACTACCTTTAGTTCGTTTAAACACG |
| qPCR rev primer 100 nt control RNA | CATGATCAAATTGACCCAAAGTTTC |
| qPCR fwd primer GcvB | GCCGGAACGAAAAGTTTTATCGG |
| qPCR rev primer GcvB | TGCTACCATGGTCTGAATCGC |
| qPCR fwd primer GlmY | GTGGCTCATTCACCGACTTATG |
| qPCR rev primer GlmY | CCCGATGGTTGATATAGCTACG |
| qPCR fwd primer ssrS | TCTCTGAGATGTTCGCAAGCG |
| qPCR rev primer ssrS | GTGTCGTCGCAGTTTTAAGGC |
| qPCR fwd primer acpP | TGAGCACTATCGAAGAACGCG |
| qPCR rev primer acpP | CTCAACGGTGTCAAGAGAATCC |

| | |
|---|---|
| qPCR fwd primer dda.1 | GGTTTATGTATATGCGATAGTTTACCG |
| qPCR rev primer dda.1 | TCTTTAAGAGTGGTAAATACTTTATCAGC |
| qPCR fwd primer motA | TGTCTAAAGTAACTTACATCATCAAAGC |
| qPCR rev primer motA | GCACCTCACGAACTTCTGC |
| qPCR fwd primer alc | TTTACAACTTATTACTACTGAAATGGTCG |
| qPCR rev primer alc | CTTAGCTAAATCTTTCTTAAGACCAGTG |
| qPCR fwd primer segG | GGTTTACATTTGAAGACCGTGTC |
| qPCR rev primer segG | CAGTATTCCTTTCCGGAAGAGTC |
| qPCR fwd primer nrdC.4 | GAATATATCAAATCATTCAATAGCG |
| qPCR rev primer nrdC.4 | CTTGCATCAACACTATTGAGCC |
| qPCR fwd primer 5S | TTGAAGAGTTTGATCATGGCTCAG |
| qPCR rev primer 5S | CAGTTTCCCAGACATTACTCACC |
| **circNC-like protocol, CapPro assay** | |
| RT primer dda.1 circNC and CapPro | CTTTTCAGTTTAAGTTTATCAATAAAAGAC |
| qPCR rev primer dda.1 circNC and CapPro | CAAATTTTGATACCGCTCTACACC |
| qPCR fwd primer dda.1 circNC and CapPro | GATATCCGCTAAATTGTTCAATTTTTAAAC |
| RT primer gcvB circNC | CAGAACACGCATTCCGATAAAAC |
| qPCR rev primer gcvB circNC | TTCGTTCCGGCTCAGGAAG |
| qPCR fwd primer gcvB circNC | TGAACTTTTGGCTTACGGTTGTG |
| RT primer modA_modB circNC | TATAGATAACTGTTCAACATCTGCAAG |
| qPCR rev primer modA modB circNC | CCAGCTGGAAGTGTATTTAGAATTG |
| qPCR fwd primer modA modB circNC | AATCATGATAGCCTCCGTATACTTC |
| RT primer NAD-ssrS circNC | ACCGAGCATGCTCACCAAC |
| qPCR rev primer NAD-ssrS circNC | GTCCGAGAAGCCTTAAAACTGC |
| qPCR fwd primer NAD-ssrS circNC | CGGAGCGCCACATTCTTG |
| RT primer RNAI circNC and CapPro | CGAAGGTAACTGGCTTCAGC |
| qPCR rev primer RNAI circNC and CapPro | AGAGCGCAGATACCAAATACTG |

| | |
|---|---|
| qPCR fwd primer RNAI circNC and CapPro | AAAAAGAGTTGGTAGCTCTTGATC |
| RT primer ssrS circNC and CapPro | AGGGGACTGGCCCGC |
| qPCR rev primer ssrS circNC and CapPro | GCGAACATCTCAGAGAAATTTTGTC |
| qPCR fwd primer ssrS circNC and CapPro | GCCGATATTTCATACCACAAGAATG |
| RT primer 5S rRNA circNC | GGTCAGGTGGGACCACC |
| qPCR rev primer 5S rRNA circNC | CGCTACTGCCGCCAGG |
| qPCR fwd primer 5S rRNA circNC | TGCCGAACTCAGAAGTGAAACG |
| **Cloning and mutagenesis of *nudE.1* gene** | |
| NudE.1 fwd NcoI | ATCGACCCATGGGACAGGAAATTAAAATGAAAACATTATCAGC |
| NudE.1 rev XhoI | GTGCTCGAGGCCCTGAAAATAAAGATTCTCACCAAAGAGGTTGTTCATTATTCG GTAAAG |
| NudE.1 E64,65Q fwd | CGAAGAGAATGTTTACAACAGACTGGTTTTAGC |
| NudE.1 E64,65Q rev | TGCTGCATCTAATGCGCTTAAATC |
| NudE.1 screening fwd | CCAGTCACGACGTTGTAAAACGCATAATACCTCCTAAGTATTTATAGAAGG |
| NudE.1 screening rev | AGCGGATAACAATTTCACACAGGCTAGGGATATGGCGTATGTTCATTGAAATGC C |
| **NAD captureSeq** | |
| Adenylated RNA 3′-adapter | rAppCNNNNNNAGATCGGAAGAGCACACGTCTG-3SpC3 |
| RT primer | CAGACGTGTGCTCTTCCGAT |
| cDNA anchor fwd | pCAGATCGGAAGAGCGTCGTGTCCC-SpC3 |
| cDNA anchor rev | ACACGACGCTCTTCCGATCTGGG |
| BC1 fwd | CACAAAGACACCGACAACTTTCTTGGGACACGACGCTCTTCCGATCTG |
| BC1 rev | CACAAAGACACCGACAACTTTCTTCAGACGTGTGCTCTTCCGAT |
| BC2 fwd | ACAGACGACTACAAACGGAATCGAGGGACACGACGCTCTTCCGATCTG |
| BC2 rev | ACAGACGACTACAAACGGAATCGACAGACGTGTGCTCTTCCGAT |
| BC3 fwd | CCTGGTAACTGGGACACAAGACTCGGGACACGACGCTCTTCCGATCTG |
| BC3 rev | CCTGGTAACTGGGACACAAGACTCCAGACGTGTGCTCTTCCGAT |
| BC4 fwd | TAGGGAAACACGATAGAATCCGAAGGGACACGACGCTCTTCCGATCTG |
| BC4 rev | TAGGGAAACACGATAGAATCCGAACAGACGTGTGCTCTTCCGAT |
| BC5 fwd | AAGGTTACACAAACCCTGGACAAGGGGACACGACGCTCTTCCGATCTG |
| BC5 rev | AAGGTTACACAAACCCTGGACAAGCAGACGTGTGCTCTTCCGAT |
| BC6 fwd | GACTACTTTCTGCCTTTGCGAGAAGGGACACGACGCTCTTCCGATCTG |
| BC6 rev | GACTACTTTCTGCCTTTGCGAGAACAGACGTGTGCTCTTCCGAT |
| BC7 fwd | AAGGATTCATTCCCACGGTAACACGGGACACGACGCTCTTCCGATCTG |
| BC7 rev | AAGGATTCATTCCCACGGTAACACCAGACGTGTGCTCTTCCGAT |
| BC8 fwd | ACGTAACTTGGTTTGTTCCCTGAAGGGACACGACGCTCTTCCGATCTG |
| BC8 rev | ACGTAACTTGGTTTGTTCCCTGAACAGACGTGTGCTCTTCCGAT |
| BC9 fwd | AACCAAGACTCGCTGTGCCTAGTTGGGACACGACGCTCTTCCGATCTG |
| BC9 rev | AACCAAGACTCGCTGTGCCTAGTTCAGACGTGTGCTCTTCCGAT |
| BC10 fwd | GAGAGGACAAAGGTTTCAACGCTTGGGACACGACGCTCTTCCGATCTG |
| BC10 rev | GAGAGGACAAAGGTTTCAACGCTTCAGACGTGTGCTCTTCCGAT |
| BC11 fwd | TCCATTCCCTCCGATAGATGAAACGGGACACGACGCTCTTCCGATCTG |

| | |
|---|---|
| BC11 rev | TCCATTCCCTCCGATAGATGAAACCAGACGTGTGCTCTTCCGAT |
| BC12 fwd | TCCGATTCTGCTTCTTTCTACCTGGGGACACGACGCTCTTCCGATCTG |
| BC12 rev | TCCGATTCTGCTTCTTTCTACCTGCAGACGTGTGCTCTTCCGAT |
| ***In vitro* transcription templates** | |
| Fwd ultramer IVT 100 nt control RNA | TAATACGACTCACTATTATCTTGATACTACCTTTAGTTCGTTTAAACACGTTCTTG ATAGTATC*TTTTTATT*AACC |
| Rev ultramer IVT 100 nt control RNA | CATGATCAAATTGACCCAAAGTTTCAACGCTTTACGCGTTGGGTTAATAAAAAG ATACTATCAAGAACGTG |
| Fwd primer IVT 100 nt control RNA | TAATACGACTCACTATTATCTTGATACTACCTTTAG |
| Rev primer IVT 100 nt control RNA | CATGATCAAATTGACCCAAAGTTTCAACGCTTTACGCG |
| Fwd ultramer IVT RNAI | TAATACGACTCACTATTACAGTATTTGGTATCTGCGCTCTGCTGAAGCCAGTTAC CTTCGGAAAAAGAGTTGGTAGCTCTTG |
| Rev ultramer IVT RNAI | AACAAAAAAACCACCGCTACCAGCGGTGGTTTGTTTGCCGGATCAAGAGCTAC CAACTCTTTTTCCGAAGGTAACTGGCTTC |
| Fwd primer IVT RNAI | TAATACGACTCACTATTACAG |
| Rev primer IVT RNAI | AACAAAAAAACCACCGCTACC |

**Supplementary Table S3: RNA sequences used in this study.**

| **Name** | **Sequence** |
|---|---|
| 100 nt control RNA | AUCUUGAUACUACCUUUAGUUCGUUUAAACACGUUCUUGAUAGUAUCUUU UUAUUAACCCAACGCGUAAAGCGUUGAAACUUUGGGUCAAUUUGAUCAUG |
| RNAI | ACAGUAUUUGGUAUCUGCGCUCUGCUGAAGCCAGUUACCUUCGGAAAAAG AGUUGGUAGCUCUUGAUCCGGCAAACAAACCACCGCUGGUAGCGGUGGUU UUUUUGUU |
| 10mer-Cy5 | pACAGUAUUUG-Cy5 |
| Linear-10mer-Cy5 | pAGACUUCGAC-Cy5 |
| 2nt-5′-overhang-10mer-Cy5 | pACAGACUUCGGUCU-Cy5 |
| Blunt-end-10mer-Cy5 | pAGACUUCGGUCU-Cy5 |
| 1nt-3′-overhang-10mer-Cy5 | pAGACUUCGGUCUA-Cy5 |

**Supplementary Table S4: DNA sequences of proteins as expressed from plasmids used in this study.** Full plasmid maps are available at: https://github.com/MaikTungsten/PhageEpitranscriptomics.

| **Plasmid Name** | **Sequence** | **Reference** |
|---|---|---|
| pET28-NadR | ATGTCGTCATTTGATTACCTGAAAACTGCCATCAA GCAACAGGGCTGCACGCTACAGCAGGTAGCTGA TGCCAGCGGTATGACCAAAGGGTATTTAAGCCAG TTACTGAATGCCAAAATCAAAAGCCCCAGCGCGC AAAAGCTGGAGGCGTTGCACCGTTTTTTGGGGCT TGAGTTTCCCCGGCAGAAGAAAACGATCGGTGTC GTATTCGGTAAGTTCTACCCACTGCATACCGGAC ATATCTACCTTATCCAGCGCGCCTGTAGCCAGGTT | This study; originally described in Höfer, K., PhD thesis, University of Heidelberg (2017) |

| | | |
|---|---|---|
| | GACGAGCTGCATATCATTATGGGTTTTGACGATA<br>CCCGTGACCGCGCGTTGTTCGAAGACAGTGCCAT<br>GTCGCAGCAGCCGACCGTGCCGGATCGTCTGCGT<br>TGGTTATTGCAAACTTTTAAATATCAGAAAAATAT<br>TCGCATTCATGCTTTCAACGAAGAGGGCATGGAG<br>CCGTATCCGCACGGCTGGGATGTGTGGAGCAAC<br>GGCATCAAAAAGTTTATGGCTGAAAAAGGGATCC<br>AGCCGGATCTGATCTACACCTCGGAAGAAGCCG<br>ATGCGCCACAGTATATGGAACATCTGGGGATCGA<br>GACGGTGCTGGTCGATCCGAAACGTACCTTTATG<br>AGTATCAGCGGTGCGCAGATCCGCGAAAACCCG<br>TTCCGCTACTGGGAATATATTCCTACCGAAGTGA<br>AGCCGTTTTTTGTGCGTACCGTGGCGATCCTTGGC<br>GGCGAGTCGAGCGGTAAATCCACCCTGGTAAAC<br>AAACTTGCCAATATCTTCAACACCACCAGTGCGT<br>GGGAATATGGCCGCGATTATGTCTTTTCACACCTC<br>GGCGGTGATGAGATCGCATTGCAGTATTCTGACT<br>ACGATAAAATCGCGCTGGGCCACGCTCAATACAT<br>TGATTTTGCGGTGAAATATGCCAATAAAGTGGCA<br>TTTATCGATACCGATTTTGTCACCACTCAGGCGTT<br>CTGCAAAAAGTACGAAGGGCGGGAACATCCGTT<br>CGTGCAGGCGCTGATTGATGAATACCGTTTCGAT<br>CTGGTGATCCTGCTGGAGAACAACACGCCGTGG<br>GTGGCGGATGGTTTACGCAGCCTCGGCAGTTCGG<br>TGGATCGCAAAGAGTTCCAGAACTTGCTGGTGG<br>AGATGCTCGAAGAGAACAATATCGAATTTGTGCG<br>GGTTGAAGAGGAAGATTACGACAGTCGTTTCCTG<br>CGCTGCGTGGAACTGGTGCGGGAGATGATGGGG<br>GAGCAGAGATAA | |
| pET28-NudE.1 WT | ATGGGACAGGAAATTAAAATGAAAACATTATCAG<br>CTGGTATTATCTTTATGACAGAAGATAAAGATTTA<br>TTTATGGGTCGGGTTACTGGTTCTCGTAAGACTGG<br>AATGATGGCACATCGTTGGGATATTCCAAAGGGC<br>CGTGTAGAAAATTCTGATTTAAGCGCATTAGATG<br>CAGCACGAAGAGAATGTTTAGAAGAGACTGGTTT<br>TAGCAATTATAATCCAGACCTTCTAGAAGACCTA<br>GGTGTATTTAAATATTCTAGTAATAAAGACCTACA<br>GTTATTTTATTACACGATTCCAGTAGAGCATGAGA<br>TGTTCAGAAATTGCCGTTGCGAGTCTTATTTTGAA<br>AATAAAGATGGCGTTATGATTCCAGAGATGGACG<br>CTTTTGCTCTTATTCCTCGTACTCAGTGGCAATATG<br>TGATGGGTCCTTCACTTTACCGAATAATGAACAAC<br>CTCTTTGGTGAGAATCTTTATTTTCAGGGCCTCGA<br>GCACCACCACCACCACCACTGA | This study |
| pET28-NudE.1<br>E64,65Q | ATGGGACAGGAAATTAAAATGAAAACATTATCAG<br>CTGGTATTATCTTTATGACAGAAGATAAAGATTTA<br>TTTATGGGTCGGGTTACTGGTTCTCGTAAGACTGG<br>AATGATGGCACATCGTTGGGATATTCCAAAGGGC<br>CGTGTAGAAAATTCTGATTTAAGCGCATTAGATG<br>CAGCACGAAGAGAATGTTTACAACAGACTGGTTT<br>TAGCAATTATAATCCAGACCTTCTAGAAGACCTA<br>GGTGTATTTAAATATTCTAGTAATAAAGACCTACA<br>GTTATTTTATTACACGATTCCAGTAGAGCATGAGA | This study |

| | | |
|---|---|---|
| | TGTTCAGAAATTGCCGTTGCGAGTCTTATTTTGAA<br>AATAAAGATGGCGTTATGATTCCAGAGATGGACG<br>CTTTTGCTCTTATTCCTCGTACTCAGTGGCAATATG<br>TGATGGGTCCTTCACTTTACCGAATAATGAACAAC<br>CTCTTTGGTGAGAATCTTTATTTTCAGGGCCTCGA<br>GCACCACCACCACCACCACTGA | |
| pET28-NudC WT | ATGGATCGTATAATTGAAAAATTAGATCACGGCT<br>GGTGGGTCGTCAGCCATGAACAAAAATTATGGTT<br>GCCGAAGGGAGAATTGCCATATGGCGAAGCGGC<br>AAATTTCGATCTTGTGGGTCAGCGCGCACTACAA<br>ATCGGCGAATGGCAGGGGGAACCTGTTTGGTTAG<br>TACAACAGCAGCGGCGTCACGATATGGGGTCGG<br>TACGTCAGGTCATTGATCTCGATGTTGGGCTGTTT<br>CAACTGGCCGGACGAGGCGTACAACTGGCGGAG<br>TTTTACCGATCGCATAAATACTGTGGTTACTGCGG<br>GCATGAAATGTATCCGAGCAAAACCGAATGGGC<br>GATGCTGTGCAGCCATTGCCGTGAGCGTTACTAC<br>CCGCAAATCGCCCCCTGCATTATTGTTGCCATCCG<br>TCGCGATGATTCGATCCTCCTCGCCCAGCATACC<br>CGCCATCGTAACGGTGTCCATACAGTACTTGCCG<br>GATTCGTCGAAGTGGGCGAAACCCTCGAGCAGG<br>CAGTCGCGCGGGAAGTGATGGAAGAGAGCGGAA<br>TTAAAGTTAAAAACTTGCGTTACGTGACTTCTCAG<br>CCGTGGCCGTTTCCTCAGTCTTTAATGACCGCGTT<br>TATGGCGGAATATGACAGCGGCGACATCGTGATC<br>GACCCGAAAGAATTGCTCGAGGCGAACTGGTATC<br>GCTATGACGATTTGCCGTTACTCCCGCCGCCCGG<br>CACCGTAGCGCGCCGTCTGATAGAAGATACGGT<br>GGCGATGTGTCGGGCAGAGTATGAGCTGGTGCC<br>GCGCGGCAGCGCGGCCGCACTCGAGCACCACCA<br>CCACCACCAC | (Höfer et al. 2016) |
| pUC19 | No insert. | (Cahova et al. 2015) |
| pCpf NudE.1 | Inserted guide RNA against NudE.1:<br>GAAGAGACTGGTTTTAGCAA | This study |
| pBA560-Cas13a-NudE.1 | Inserted guide RNA against NudE.1:<br>GCTAAAACCAGTCTCTTCTAAACATTCTCTT | This study, (Adler et al. 2022) |
| pET28 NgTET_NudE.1 E64,65Q | NgTET insert:<br>ATGGGAACGACATTTAAACAGCAGACGATTAAAG<br>AAAAAGAGACAAAGCGTAAATACTGTATCAAAG<br>GGACCACTGCGAATCTGACACAAACCCATCCCAA<br>TGGGCCAGTGTGTGTTAACCGCGGGGAGGAAGT<br>AGCAAATACGACTACTCTGTTGGACTCAGGGGGC<br>GGGATTAACAAAAAATCGCTGTTGCAGAATCTGT<br>TGTCCAAATGTAAAACTACATTTCAGCAGTCATTT<br>ACAAACGCCAACATTACTTTAAAGGATGAAAAGT<br>GGCTTAAAAACGTCCGTACTGCTTATTTTGTTTGC<br>GATCATGACGGGAGCGTCGAACTTGCATACCTTC<br>CTAACGTGCTTCCCAAAGAATTAGTCGAAGAATT<br>CACCGAGAAATTTGAATCGATCCAGACCGGACGT<br>AAGAAAGACACAGGTTACTCAGGTATTCTGGACA<br>ACTCGATGCCGTTCAATTACGTCACTGCGGATTTA<br>TCACAGGAGTTAGGACAGTATCTGTCTGAGATTG<br>TGAATCCTCAGATCAACTATTACATCAGTAAATTG | This study, (Pozhydaieva et al. 2024a; Pozhydaieva et al. 2024b) |

| | |
|---|---|
| | CTGACTTGCGTTAGTTCACGTACAATCAATTACCT GGTATCTTTGAATGACTCGTACTATGCCCTTAACA ACTGTTTGTATCCTTCAACCGCCTTCAACTCATTA AAGCCGTCCAACGACGGCCACCGCATCCGTAAA CCTCATAAGGACAATTTGGACATTACCCCGTCGA GCCTTTTCTATTTTGGAAATTTTCAAAATACGGAA GGATATCTTGAGTTAACAGACAAGAATTGCAAAG TTTTCGTCCAACCGGGGGATGTATTATTTTTCAAA GGCAATGAATATAAACACGTCGTGGCCAATATCA CCTCGGGCTGGCGCATTGGATTGGTCTACTTCGC ACACAAGGGGAGTAAAACTAAACCGTATTATGAA GACACGCAGAAGAACTCCCTGAAAATTCATAAAG AAACGAAATAAC<br>NudE.1 E64,65Q insert:<br>GACAGGAAATTAAAATGAAAACATTATCAGCTGG TATTATCTTTATGACAGAAGATAAAGATTTATTTAT GGGTCGGGTTACTGGTTCTCGTAAGACTGGAATG ATGGCACATCGTTGGGATATTCCAAAGGGCCGTG TAGAAAATTCTGATTTAAGCGCATTAGATGCAGC ACGAAGAGAATGTTTACAACAGACTGGTTTTAGC AATTATAATCCAGACCTTCTAGAAGACCTAGGTG TATTTAAATATTCTAGTAATAAAGACCTACAGTTA TTTTATTACACGATTCCAGTAGAGCATGAGATGTT CAGAAATTGCCGTTGCGAGTCTTATTTTGAAAATA AAGATGGCGTTATGATTCCAGAGATGGACGCTTT TGCTCTTATTCCTCGTACTCAGTGGCAATATGTGA TGGGTCCTTCACTTTACCGAATAATGAACAACCTC TTT |

**Supplementary Table S5: Mass transitions, collision energies, cell accelerator voltages and dwell times for LC-MS/MS analysis of metabolites NAD, FAD, UDP-GlcNac.** Parameters have been optimized using chemically pure standards.

| Name | Precurser Ion | Product Ion | Collision energy [V] | Fragmentor Voltage [V] | Cell Accelorator Voltage [V] | Dwell time [msec] | Polarity |
|---|---|---|---|---|---|---|---|
| NAD | 664.1 | 524<br>428 | 18<br>26 | 380 | 5 | 90 | Positive |
| FAD | 786.2 | 348.1<br>136 | 21<br>46 | 380 | 5 | 90 | Positive |
| UDP-GlcNac | 606.1 | 384.8<br>281.90 | 29<br>32 | 380 | 5 | 90 | Negative |

### Captions for separate Supplementary Tables

**Supplementary Table S6: NAD-RNAs identified by NAD captureSeq.**
Data is presented for replicate 1 (**A**) and replicate 2 (**B**). For each time point, it is indicated, whether the transcript of the corresponding gene was found enriched (+) or not (-). Entity represents the species, which is either *E. coli* (U00096.3), T4 phage (NC_000866.4) or the 100 nt control RNA (spike-in).

**Supplementary Table S7: qPCR to confirm enrichment of NAD-RNAs on cDNA level as reported by NAD captureSeq.**
For each analyzed time point and target gene/cDNA the Ct-value of technical duplicates is presented. The Log2 Fold Change (LFC) is calculated as the difference of Ct-values for –ADPRC and +ADPRC sample. Negative LFCs are marked in red, ct (no template) indicates background signal. Both *E. coli* and T4 phage targets have been validated.

**Supplementary Table S8: Statistics of Transcription Start Site Prediction for *E. coli* (A) and T4 phage (B).**

**Supplementary Table S9: Burst size of T4 phage WT and NudE.1 M1 mutant as a mean of technical triplicates.**

**Supplementary Table S10: Top 100 hits of protein blast search of NudE.1 protein sequence (Y06L).**
